# Nucleosome Positioning Shapes Cryptic Antisense Transcription

**DOI:** 10.1101/2025.07.14.664753

**Authors:** Jian Yi Kok, Zachary H. Harvey, Elin Axelsson, Frédéric Berger

## Abstract

Maintaining transcriptional fidelity is essential for precise gene regulation and genome stability. Despite this, cryptic antisense transcription, occurring opposite to canonical coding sequences, is a pervasive feature across all domains of life. How such potentially harmful cryptic sites are regulated remains incompletely understood. Here, we show that nucleosome arrays within gene bodies play a key role in suppressing cryptic transcription. Using the fission yeast *Schizosaccharomyces pombe* as a model, we demonstrate that CHD1-family chromatin remodelers coordinate with the transcription elongation machinery, specifically the PAF complex, to position nucleosomes at sites of cryptic transcription initiation within gene bodies. In the absence of CHD1, AT-rich sequences within gene bodies lose nucleosome occupancy, exposing promoter-like sequences that drive cryptic initiation. While cryptic transcription is generally detrimental, we identify a subset of antisense transcripts that encode critical meiotic genes, suggesting that cryptic transcription can also serve as a source of regulatory innovation. These findings underscore the essential role of nucleosome remodelers in maintaining transcriptional fidelity and reveal their broader contributions to cellular homeostasis and evolutionary adaptability.

## INTRODUCTION

Antisense transcription, a pervasive phenomenon with deep evolutionary roots^1^, occurs when RNA polymerase II (RNAPII) initiates transcription in the opposite direction of protein-coding genes^2^. Observed across prokaryotes^3–5^, unicellular protists^6^, plants^7^ and animals^8,9^, antisense transcription has been implicated in diverse regulatory functions, including the modulation of sense transcript levels^8,10–12^, epigenetic regulation^13^, and developmental processes^14–16^. However, when misregulated, it can lead to transcriptional interference^17^ and genome instability^18^, fueling the pathology of cancers and neurological disorders^19–22^. Despite its widespread occurrence, how cells balance the regulatory roles of antisense transcripts with the need to suppress their potentially harmful effects when misregulated remains poorly understood.

Chromatin plays an important part in transcription start site (TSS) selection, and in transcriptional regulation. Nucleosomes—the basic units of chromatin composed of two subunits each of the histones H2A, H2B, H3 and H4 wrapped by ∼147 bp of DNA—act as both physical barriers and dynamic regulators of transcription across the transcriptional unit^23–25^. Beyond the well-defined roles of the +1 nucleosome at the TSS^26–28^, nucleosomes are also regularly positioned and spaced along gene bodies, forming stereotypical phased arrays^29,30^. Although nucleosome arrays have been proposed to play roles in transcriptional elongation and termination^31^, preventing spurious transcription initiation^32–34^ and safeguarding the genome^35^, their role in regulating sense transcription is modest, and therefore the precise link between their placement and gene regulation has remained a subject of debate.

Mechanistic studies seeking to address the role of nucleosome arrays in transcription have predominantly focused on SWI/SNF nucleosome remodelers, particularly CHD1 and ISWI^36–39^, which are broadly essential across eukaryotes and are known to play important roles both in maintaining chromatin structure and regulating transcription^40–43^. Whereas their role in this context has been proposed, in animals and plants, multiple paralogs of ISWI and CHD1 have evolved^44–46^, making it challenging to perform efficient genetic interventions to study nucleosome phasing, and to deconvolve direct and indirect effects. For example, even in the budding yeast *Saccharomyces cerevisiae*, the functions of CHD1 and the two ISWI paralogs, ISW1 and ISW2, appear to overlap, contributing redundantly to proper nucleosome phasing and transcriptional termination^31,35,42^. Additionally, the INO80 remodeling complex also plays a role in nucleosome spacing^47–50^, further complicating the analysis of phasing.

While many studies on nucleosome remodelers have focused on the action of their conserved ATPase domain which drives nucleosome remodeling^51–53^, the roles of their accessory domains remain an area of active investigation. These accessory domains are increasingly recognized for their importance in targeting remodelers to specific genomic regions and coordinating interactions. For example, the chromodomains in CHD1-family remodelers bind methylated H3 tails^54^ for proper localization, while the HAND-SANT-SLIDE domain in ISWI-family remodelers is critical for recognizing linker DNA, which helps space nucleosomes^55^. Despite these insights, the precise contributions of several other accessory domains to overall remodeler function and transcriptional regulation are still unclear.

Here, we leverage the fission yeast *Schizosaccharomyces pombe,* which lacks ISWI-family remodelers^56^, placing greater reliance on CHD1-family remodelers to maintain gene body nucleosome organization. This unique feature makes *S. pombe* an ideal model to isolate and study the specific roles of remodelers in nucleosome phasing and transcriptional regulation. Using transcriptomics, chromatin profiling, and functional assays, we show that the CHD1-family remodeler Hrp3 in fission yeast establishes nucleosome arrays within gene bodies, counteracting cryptic antisense initiation. We further demonstrate that Hrp3 interacts with Prf1, a subunit of the Paf1 elongation complex, via its CHCT domain, linking nucleosome remodeling to transcription elongation. Loss of Hrp3 or disruption of its interaction with Prf1 results in impaired nucleosome phasing, increased antisense transcription, and significant fitness defects under stress conditions. Additionally, we identify thousands of antisense transcripts, including functional protein-coding transcripts and transcripts with unknown coding potential. Many of these are in fact sense transcripts of genes nested within larger host genes in a convergent orientation, which depend on Hrp3 for their regulation. These findings provide new insights into the mechanisms by which nucleosome remodelers suppress cryptic transcription and coordinate expression of nested genes, offering a framework to better understand how their dysregulation may contribute to altered gene expression programs and disease.

## RESULTS

### Hrp3 is the Predominant SWI/SNF Remodeler Regulating Nucleosome Phasing and Antisense Transcription

To systematically investigate the functional impact of nucleosome remodelers on antisense transcription in the fission yeast *Schizosaccharomyces pombe*, we generated knockout mutants for 11 viable nucleosome remodelers. This included three groups of paralogs - the FUN30 group (*fft1*, *fft2*, and *fft3*), the CHD1 group (*hrp1* and *hrp3*), and the ULS1 group (*rrp1* and *rrp2*), as well as four additional remodelers: i*rc20*, *mit1*, *snf22*, and *swr1*, the latter being involved in the exchange of the transcription-associated histone variant H2A.Z^57^ (**Fig. 1A**). Since remodelers are primarily associated with nucleosome dynamics, we assessed the impact of these mutations by first measuring genome-wide changes in nucleosome organization, followed by correlating these changes with sense and antisense transcription.

**Figure 1.**
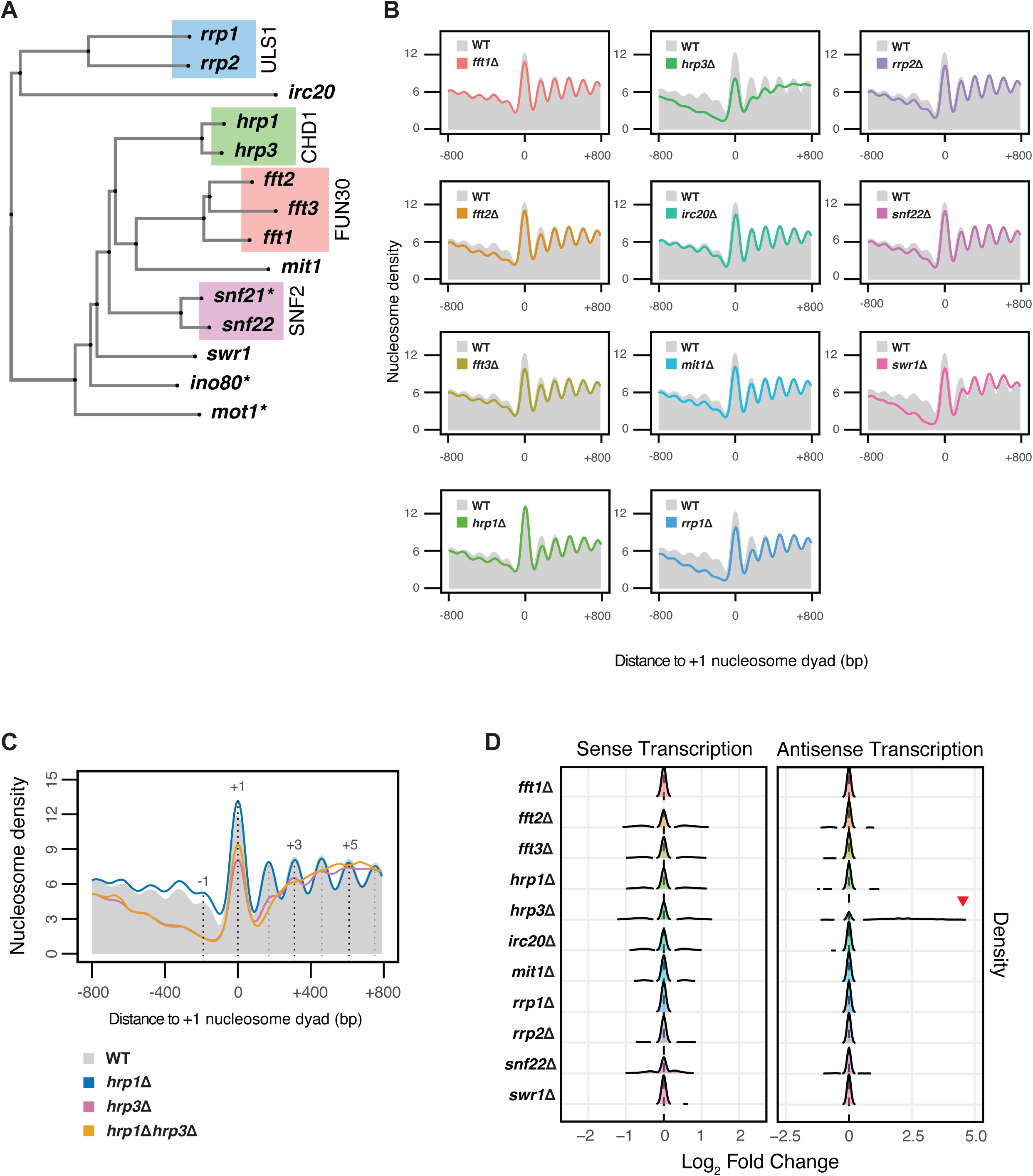
The Impact of Nucleosome Remodelers on Nucleosome Phasing and Transcription. (A) Unrooted tree generated from multiple sequence alignment of 14 nucleosome remodelers in *S. pombe*. Groupings of paralogs are indicated by coloring, with each group corresponding to the ortholog of a nucleosome remodeler in *S. cerevisiae*. Asterisks (*) indicate the remodelers that are essential in *S. pombe*. (B) Metaplots of normalized nucleosome density from MNase-seq centered on the +1 nucleosome dyad for *fft1*Δ, *fft2*Δ, and *fft3*Δ, *hrp1*Δ, *hrp3*Δ, *irc20*Δ, *mit1*Δ, *rrp1*Δ, *rrp2*Δ, *snf22*Δ, and *swr1*Δ mutants. Profiles include 800 bp upstream and downstream of the +1 nucleosome dyad for all protein-coding genes. The WT profile is shown in solid grey. The plotted data represents the average signal from two biological replicates. (C) Metaplots of normalized nucleosome density from MNase-seq centered on the +1 nucleosome dyad for WT, *hrp1*Δ, *hrp3*Δ, and *hrp1*Δ*hrp3*Δ. The WT nucleosome profile is shown in solid grey, with dotted lines indicating the positions of select nucleosome maxima (- 1, +1, +3 and +5). Profiles include 800 bp upstream and downstream of the +1 nucleosome dyad for all protein-coding genes. The plotted data represents the average signal from two biological replicates. (D) Differential expression analysis of mRNA-seq data showing the distribution of log_2_fold-change values of all *S. pombe* protein-coding genes in both sense and antisense orientations across *fft1*Δ, *fft2*Δ, and *fft3*Δ, *hrp1*Δ, *hrp3*Δ, *irc20*Δ, *mit1*Δ, *rrp1*Δ, *rrp2*Δ, *snf22*Δ, and *swr1*Δ mutants compared to WT. The red arrowhead indicates the extent of antisense transcription in the *hrp3*Δ mutant. Data are calculated from three biological replicates.

To examine nucleosome organization, we performed MNase-seq to directly assess nucleosome phasing and profiled H2A.Z genomic distribution using ChIP-seq. While nucleosome arrays in wild-type (WT) and most remodeler mutant cells were highly ordered and regular (**Fig. 1B**), the *hrp3* knockout (*hrp3*11) displayed highly disrupted nucleosome phasing over gene bodies (**Fig. 1C**). Additionally, while its paralog *hrp1* (*hrp1*11) showed minimal changes to nucleosome organization, the disruption seen in *hrp3*11 was also observed in the *hrp1*11*hrp3*11 double mutant (**Fig. 1C**). Despite the strong impact of Hrp3 on nucleosome phasing, we observed no effect on H2A.Z enrichment profile in *hrp3*11 mutants (**Fig. S1A, S1B**)^58,59^, consistent with previous reports that this activity is exclusive to Swr1^57,60^. These results establish that Hrp3 is the primary CHD1-family remodeler responsible for maintaining nucleosome organization over gene bodies.

We next investigated whether these changes in nucleosome organization were associated with changes in transcription. Transcriptome analysis using RNA-seq revealed only mild changes in sense and antisense transcript levels across most mutants compared to WT, except for *hrp3*11 (**Fig. 1D**). Consistent with its strong impact on nucleosome phasing, *hrp3*11 cells exhibited up to 32-fold increase in antisense transcript levels compared to WT (**Fig. 1D**). In contrast, *hrp1*11 showed only a slight increase, consistent with previous studies^61–63^. Furthermore, *hrp1*11*hrp3*11 double mutants had antisense transcript levels comparable to those in *hrp3*11 alone (**Fig. 2A, 2B**), suggesting limited or no functional redundancy between the two paralogs. Importantly, loss of one CHD1 paralog did not trigger compensatory upregulation of the other (**Fig. S2A**). Additionally, no meaningful correlation between sense and antisense transcript levels was observed in both *hrp1Δhrp3Δ* and WT cells **(Fig. S2B–F**), suggesting that antisense transcription minimally impacts sense transcription. Together, these data demonstrate that CHD1 orthologues play a dominant role in regulating antisense transcription, with Hrp3 acting as the primary SWI/SNF remodeler responsible for suppressing antisense transcription.

**Figure 2.**
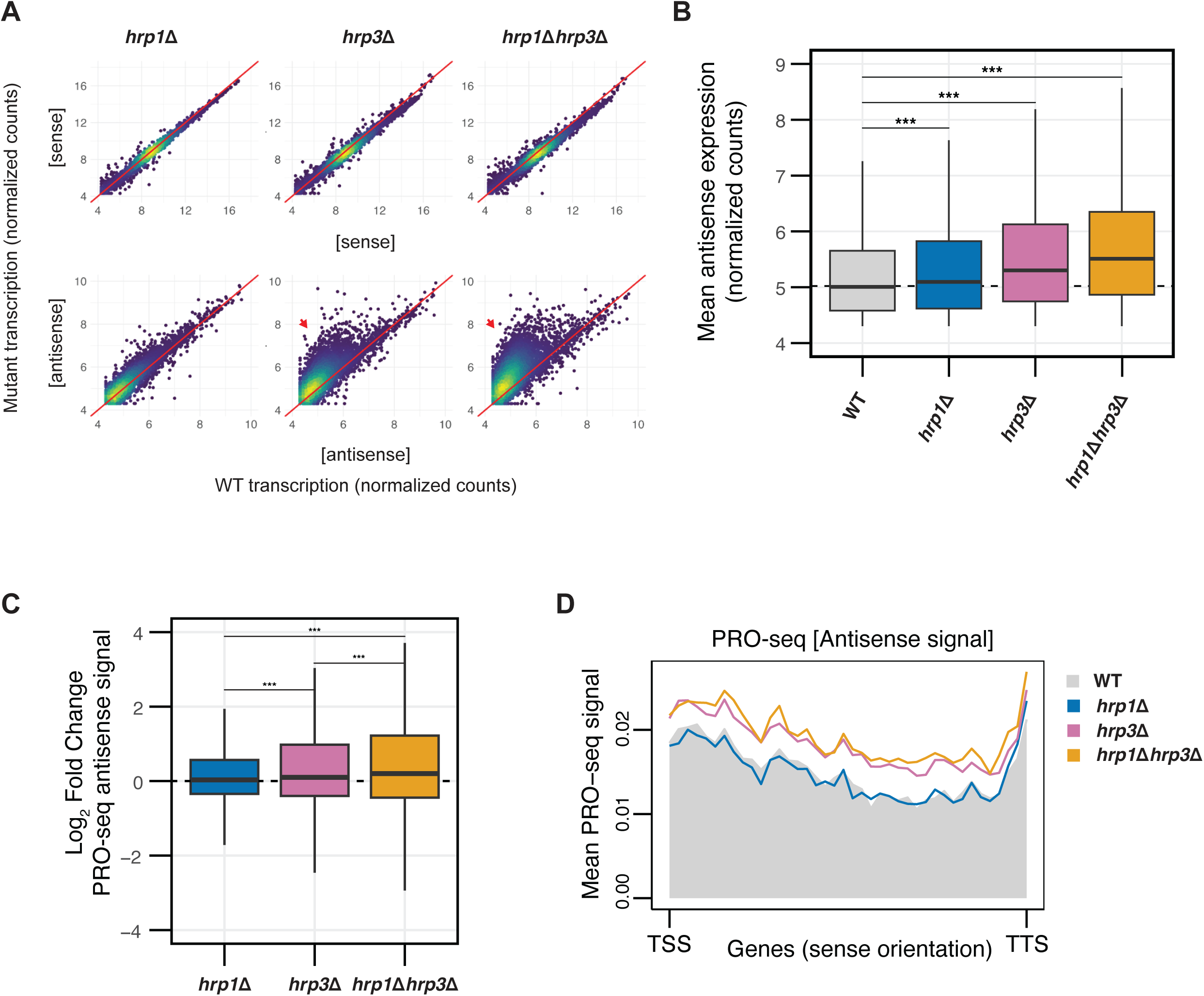
Hrp3 is the Predominant SWI/SNF Remodeler Regulating Nucleosome Phasing and Antisense Transcription. (A) Density-colored scatterplots comparing mutant (*hrp1*Δ*, hrp3*Δ *or hrp1*Δ*hrp3*Δ) transcription to WT for sense transcripts (top row) and antisense transcripts (bottom row) for all protein-coding genes. Count data is derived from variance-stabilizing transformation (vst) of raw mRNA-seq counts. Red arrows indicate elevated antisense transcription in mutants relative to WT. (B) Boxplot of normalized antisense expression counts for all antisense genes for WT, *hrp1*Δ*, hrp3*Δ and *hrp1*Δ*hrp3*Δ. The dotted line represents the median antisense expression in WT. Statistical analysis was performed using ANOVA with Tukey’s Honestly Significant Difference (HSD) test. Asterisks indicate statistical significance: p < 0.05 (*), p < 0.01 (**), p < 0.001 (***), n.s. (not significant). (C) Boxplot of log_2_fold-change values of antisense expression measured by PRO-seq for *hrp1*Δ*, hrp3*Δ and *hrp1*Δ*hrp3*Δ relative to WT. The dotted line represents a log_2_fold-change value of 0. Statistical analysis was performed using ANOVA with Tukey’s HSD test. Asterisks indicate statistical significance: p < 0.05 (*), p < 0.01 (**), p < 0.001 (***), n.s. (not significant). Data are calculated from two biological replicates. (D) Metaplots of mean antisense PRO-seq signal for *hrp1*Δ, *hrp3*Δ, and *hrp1*Δ*hrp3*Δ versus WT, plotted relative to the sense orientation of the transcription start sites (TSS) and transcription termination sites (TTS) of all genes.

### Hrp3 Blocks Cryptic Promoters to Suppress Antisense Transcription

To determine whether increased antisense transcription in *hrp3*11 reflected elevated RNA polymerase II (RNAPII) activity, we performed Precision Run-On sequencing (PRO-seq)^64^, a method that assesses nascent transcription. Consistent with the transcriptome data, *hrp3*11 and *hrp1*11*hrp3*11 mutants displayed a pronounced increase in antisense PRO-seq signal compared to *hrp1*11 across genes (**Fig. 2C, 2D),** demonstrating that the antisense expression phenotype is driven by RNAPII-dependent transcription.

We hypothesized that the increased antisense expression in *hrp3*11 was due to the shifts in the nucleosome profile that exposed regions normally occluded by nucleosomes, thereby creating permissive sites for RNAPII initiation. To test this, we first generated a high-resolution annotation of antisense transcripts in the CHD1 mutants. We used PacBio Iso-Seq^65^ to produce a comprehensive long-read transcriptome for both WT and *hrp1*11*hrp3*11. This approach yielded over 15 million high-fidelity (HiFi) reads (Q ≥ 20) with a median read length of 1,702 bp, closely matching the median transcript length in *S. pombe* (1,883 bp). Through transcript clustering and collapsing, we identified 5,105 new antisense transcripts. This more than doubles the known number of antisense transcripts, bringing the total to 9,016. Using this expanded antisense transcript annotation, we found that 4,678 antisense transcripts were differentially expressed in *hrp1Δhrp3Δ* compared to WT, overlapping with 54.9% of all protein-coding genes in *S. pombe* (**Fig. S3A**).

We next validated our new antisense transcription annotations. As a control, aligning the transcription start sites (TSS) of protein coding genes with PRO-seq profiles revealed a peak corresponding to paused or recently initiated RNAPII within 200 bp downstream of the TSS on the sense strand (**Fig. S3B**). Similarly, alignment of antisense transcript start sites (As-TSS) with PRO-seq data showed a peak immediately downstream of the As-TSS (**Fig. 3A**), validating the high accuracy and precision of our antisense transcript annotation. Consistent with the transcriptomics data, this PRO-seq peak was significantly stronger in *hrp3*11 and *hrp1*11*hrp3*11 compared to *hrp1*11 and WT (**Fig. 3A, 3B**). We noticed that these antisense peaks coincided with nucleosome-shifted regions within gene bodies, which in WT were part of a regularly phased array (**Fig. 3C**). Genes with greater nucleosome disruption in *hrp1*11*hrp3*11 exhibited higher antisense transcription (**Fig. S3C**), supporting a direct link between nucleosome disorganization and cryptic antisense expression.

**Figure 3.**
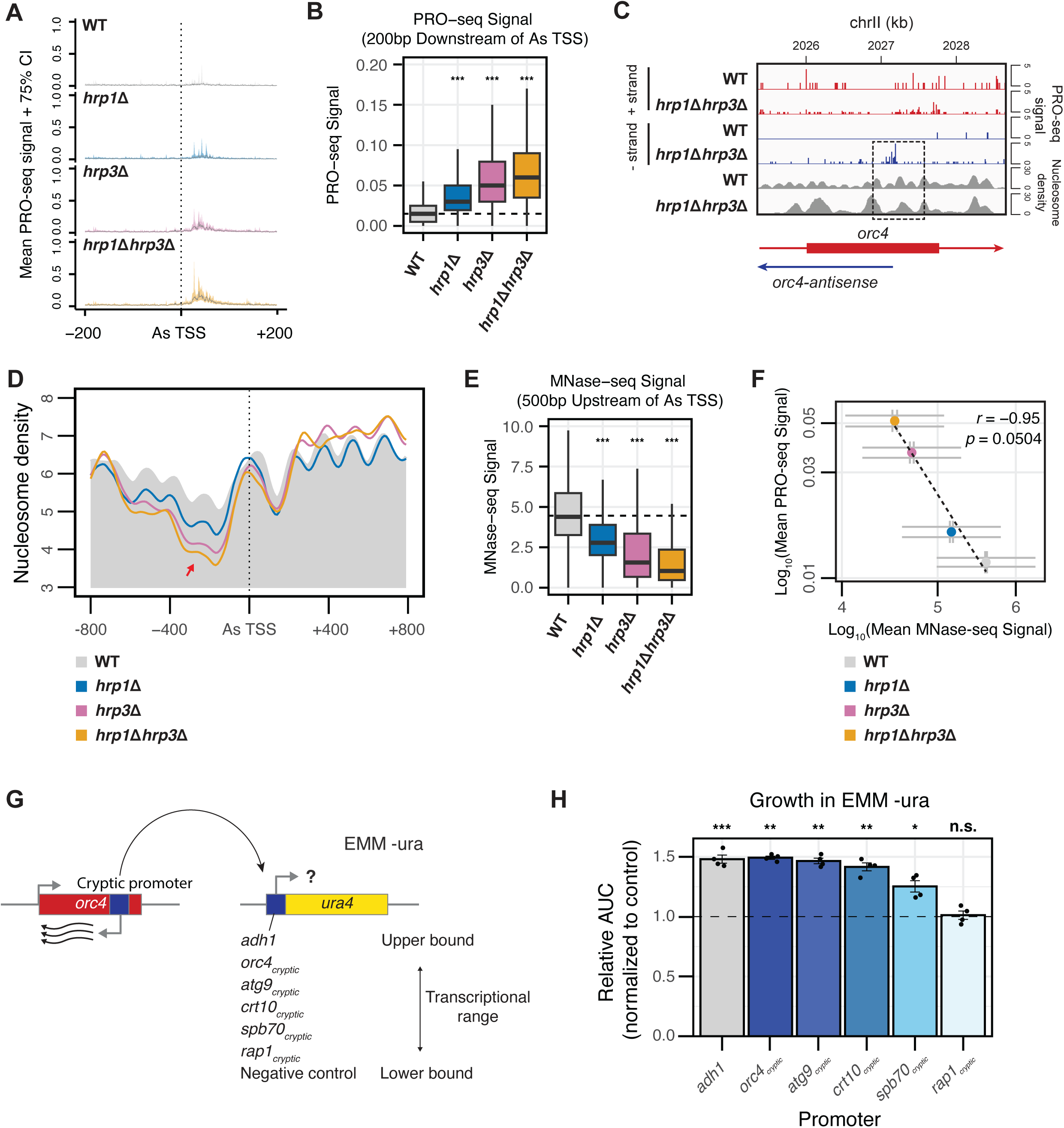
Hrp3 Blocks Cryptic Promoters to Suppress Antisense Transcription. (A) Metagene representation of mean PRO-seq signals (solid lines) with 75% confidence intervals (shaded regions) for WT, *hrp1*Δ, *hrp3*Δ, and *hrp1*Δ*hrp3*Δ mutants across the antisense (As) TSS of 4,678 differentially expressed antisense transcripts. Profiles include 200 bp upstream and downstream of the As-TSS. Data represent the mean signal from two biological replicates. (B) Boxplot of the mean PRO-seq signal in the 200 bp region downstream of the As-TSS of 4,678 differentially expressed antisense transcripts in WT, *hrp1*Δ, *hrp3*Δ, and *hrp1*Δ*hrp3*Δ. The dotted line represents the median PRO-seq signal in WT. Statistical analysis was performed using ANOVA with Tukey’s Honestly Significant Difference (HSD) test. Asterisks indicate statistical significance: p < 0.05 (*), p < 0.01 (**), p < 0.001 (***), n.s. (not significant). (C) Genome browser track of PRO-seq and MNase-seq data for the *orc4* gene, representative of genes with increased antisense transcription in *hrp1*Δ*hrp3*Δ mutants. PRO-seq signals for the + strand (red) and – strand (blue) are displayed. The dotted box highlights the region where antisense transcription initiates in the *hrp1Δhrp3Δ* mutant, coinciding with disrupted nucleosome positioning. (D) Metaplots of normalized nucleosome density from MNase-seq centered on the As-TSS of 4,678 differentially expressed antisense transcripts in WT, *hrp1*Δ, *hrp3*Δ, and *hrp1*Δ*hrp3*Δ. The WT profile is shown in solid grey. The red arrow indicates a loss of nucleosome occupancy preceding the As-TSS. Profiles include 800 bp upstream and downstream of the As-TSS. (E) Boxplot of the mean MNase-seq signal in the 500 bp region upstream of the As-TSS of 4,678 differentially expressed antisense transcripts in WT, *hrp1*Δ, *hrp3*Δ, and *hrp1*Δ*hrp3*Δ. The dotted line represents the median MNase-seq signal in WT. Statistical analysis was performed using ANOVA with Tukey’s Honestly Significant Difference (HSD) test. Asterisks indicate statistical significance: p < 0.05 (*), p < 0.01 (**), p < 0.001 (***), n.s. (not significant). (F) Scatterplot of mean PRO-seq signal in the 200 bp region downstream of the antisense TSS versus mean MNase-seq signal in the 500 bp region upstream of the antisense TSS for WT, *hrp1*Δ, *hrp3*Δ, and *hrp1*Δ*hrp3*Δ. Data is plotted for all 4,678 differentially expressed antisense transcripts between WT and *hrp1*Δ*hrp3*Δ. Spearman’s correlation and linear regression analysis were performed. Error bars represent the standard error of the mean (SEM). (G) Schematic of the *ura4*+ reporter system, where the *ura4* promoter is replaced with antisense transcription promoters from five different genes, alongside the normal *adh1* promoter (positive control) and a transcriptionally inactive exogenous sequence (negative control). Constructs were tested for their ability to drive *ura4* expression, enabling growth in EMM-ura media. (H) Bar plot showing the growth assay results for the reporter strains. The area under the curve (AUC) values from growth curves in EMM-ura media are plotted relative to the negative control. The dotted line indicates a relative AUC value of 1. Individual data points (black dots) represent biological replicates. Strains with relative AUC values above 1.0 exhibit improved growth compared to the negative control, while those below 1.0 show reduced growth. The positive control used was the *S. pombe adh1* promoter, while the negative control was a transcriptionally inactive exogenous sequence. Statistical significance was determined using pairwise t-tests comparing each strain to the negative control, with p-values adjusted using the Bonferroni method. Asterisks indicate statistical significance: p < 0.05 (*), p < 0.01 (**), p < 0.001 (***), n.s. (not significant).

To systematically investigate the relationship between nucleosome disruption and antisense initiation, we aligned the As-TSS with MNase-seq data. In all strains, a well-positioned nucleosome was observed directly over the As-TSS (**Fig. 3D**). However, in the mutants, this nucleosome was preceded by a nucleosome-depleted region, forming a pattern that closely resembled the nucleosome organization at the sense TSS. In the 500 bp region upstream of the As-TSS, *hrp3*11 and *hrp1*11*hrp3*11 mutants exhibited significant loss of nucleosome phasing and occupancy (**Fig. 3D, 3E**). In contrast, *hrp1*Δ mutants showed a smaller reduction in nucleosome occupancy in this region, accompanied by only a modest increase in antisense transcription. This suggests that the reduction in nucleosome occupancy and phasing in *hrp1*Δ alone is insufficient to induce the higher levels of antisense transcription observed in *hrp3*Δ or *hrp1*Δ*hrp3*Δ. Furthermore, a strong inverse correlation (r = −0.95) was observed between the decrease in nucleosome occupancy in this region and the antisense PRO-seq peak in each mutant (**Fig. 3F**), indicating that disrupted nucleosome organization is strongly associated with increased antisense transcription. These results suggest that, even in WT, cryptic As-TSSs are marked by nucleosomes that are more well-positioned compared to the surrounding nucleosomes. However, in the mutants, the nucleosome-depleted region immediately upstream is far more exaggerated and pronounced, likely contributing to the increased exposure of these cryptic promoter regions and the elevated levels of antisense transcription. Additionally, these findings suggest that while Hrp1 plays a minor role in maintaining nucleosome positioning, Hrp3 is the major remodeler responsible for actively positioning nucleosomes to block cryptic promoters and suppress spurious antisense initiation.

The *hrp3*Δ*hrp1*Δ mutant enabled us to investigate how nucleosomes are redistributed in the absence of extrinsic phasing activity. Previous studies have identified genomic DNA composition as a major predictor of nucleosome occupancy and binding site preference^66–68^. Consistent with a previous report^69^, we observed that nucleosomes in WT *S. pombe* align with regions of higher AT content (**Fig. S4A**). However, in *hrp1*Δ*hrp3*Δ mutants, nucleosome dyad positioning became less enriched over AT-rich regions, with an increase in the mean GC content of the associated DNA from 37.8% to 42.8% (**Fig. S4B**). Aligning the altered nucleosome profile with GC-content metaplots centered on the As-TSS revealed that, while the well-positioned nucleosome over the As-TSS in WT correlates with an AT-rich region, this nucleosome shifts toward the adjacent GC peak in all mutants (**Fig. S4C**). Furthermore, the loss of nucleosome occupancy in the upstream region in mutants is also associated with an AT-rich region. Together, these results suggest that, in the absence of Hrp3, AT-rich regions within genes become increasingly nucleosome-depleted, thereby creating nucleosome-free regions (NFRs) that we expect to contain promoters facilitating antisense transcription initiation.

To test whether these regions could act as functional promoters, we constructed a transcriptional reporter by replacing the promoter of the uracil biosynthesis gene *ura4* with 500-bp regions corresponding to the antisense promoter region from five antisense transcripts upregulated in *hrp3*Δ mutants (**Fig. 3G**). Our results showed that the *adh1* promoter used as control and four of the five 500 bp regions successfully promoted cell growth through *ura4* transcription relative to the negative control (**Fig. 3H**). Notably, the transcriptional output of these regions was comparable to that of the *adh1* promoter, supporting their intrinsic promoter activity.

These findings suggest that Hrp3 positions nucleosomes over genomic regions prone to nucleosome loss, thereby silencing otherwise antisense initiation permissive sites within genes. This conclusion is further supported by three key observations. First, 87% (4,053/4,678) of antisense transcripts initiate within protein-coding genes, while the remaining 13% initiate in nearby intergenic regions and extend into gene bodies, indicating that antisense initiation in *hrp3*Δ requires DNA sequences or nucleosome arrangements specific to gene bodies. Second, these antisense transcripts are distributed homogeneously across all chromosomes (**Fig. S3D**), suggesting that regions permissive to antisense initiation are cryptically embedded throughout most genes in the genome. Finally, antisense transcripts are also more likely to originate within longer genes (**Fig. S3E**). Taken together, these findings show that Hrp3 is essential for maintaining gene body nucleosome phasing and suppressing pervasive cryptic antisense transcription.

### Hrp3-Mediated Antisense Transcription Regulates Meiotic and Nested Gene Expression in *S. pombe*

Given that many antisense transcripts were driven by promoter-like sequences, and that natural antisense transcripts can serve diverse regulatory functions across species^10,12,70^, we next asked whether Hrp3-regulated antisense transcripts might have been co-opted for specific gene regulatory programs. We performed K-means clustering of sense genes upregulated in *hrp3*11 mutants compared to WT and mutants of other remodelers (**Fig. S5A**). This revealed a distinct group (cluster 4) enriched for genes involved in sexual reproduction (GO:0019953), whereas other clusters were depleted for these genes (**Fig. S5B**). Cross-referencing genes upregulated in *hrp3Δ* with a published list of meiotically upregulated genes (MUGs)^71^ revealed a significant overlap (**Fig. 4A, 4B**). These genes were involved in key meiotic processes including sporulation (*spo6*), septum formation (*spn6*), and meiotic transcription initiation (*atf21*) (**Table S1**). Notably, *hrp3* expression in WT cells was lowest five hours post-meiotic induction, coinciding with peak MUG expression (**Fig. 4C**), suggesting that Hrp3 normally represses meiotic gene expression during vegetative growth. A notable feature of the MUGs upregulated in *hrp3Δ* was their frequent occurrence as nested genes in a convergent orientation—completely enclosed within, and transcribed opposite to, larger host genes. Examples include *spo6*, *mfr1*, and *meu27*, which were nested within the transcripts of *lam2*, *tfg2*, and *chk1*, respectively (**Fig. 4D**, **S5C**, **S5D**). Genome-wide, we identified 212 protein coding genes in *S. pombe* with this convergent nested arrangement. To systematically explore this relationship, we performed an overlap analysis using a hypergeometric test to compare MUGs, all convergent nested genes, and genes upregulated in *hrp3*Δ (**Fig. 4E**). This analysis revealed significant pairwise overlap between all three groups, as well as a significant overlap among all three simultaneously. Volcano plots further showed that genes upregulated in *hrp3Δ* include both nested genes and nested MUGs (**Fig. 4F**). Collectively, these results indicate that Hrp3-mediated antisense transcription regulation has been co-opted to regulate the expression of nested and meiotic genes, providing an additional regulatory layer that coordinates gene expression in a context-dependent manner. In absence of Hrp3, the abundance of transcripts from non-meiotic nested genes and other antisense transcripts suggest these may either exert deleterious effects or possess functions that remain to be identified under various growth conditions.

**Figure 4.**
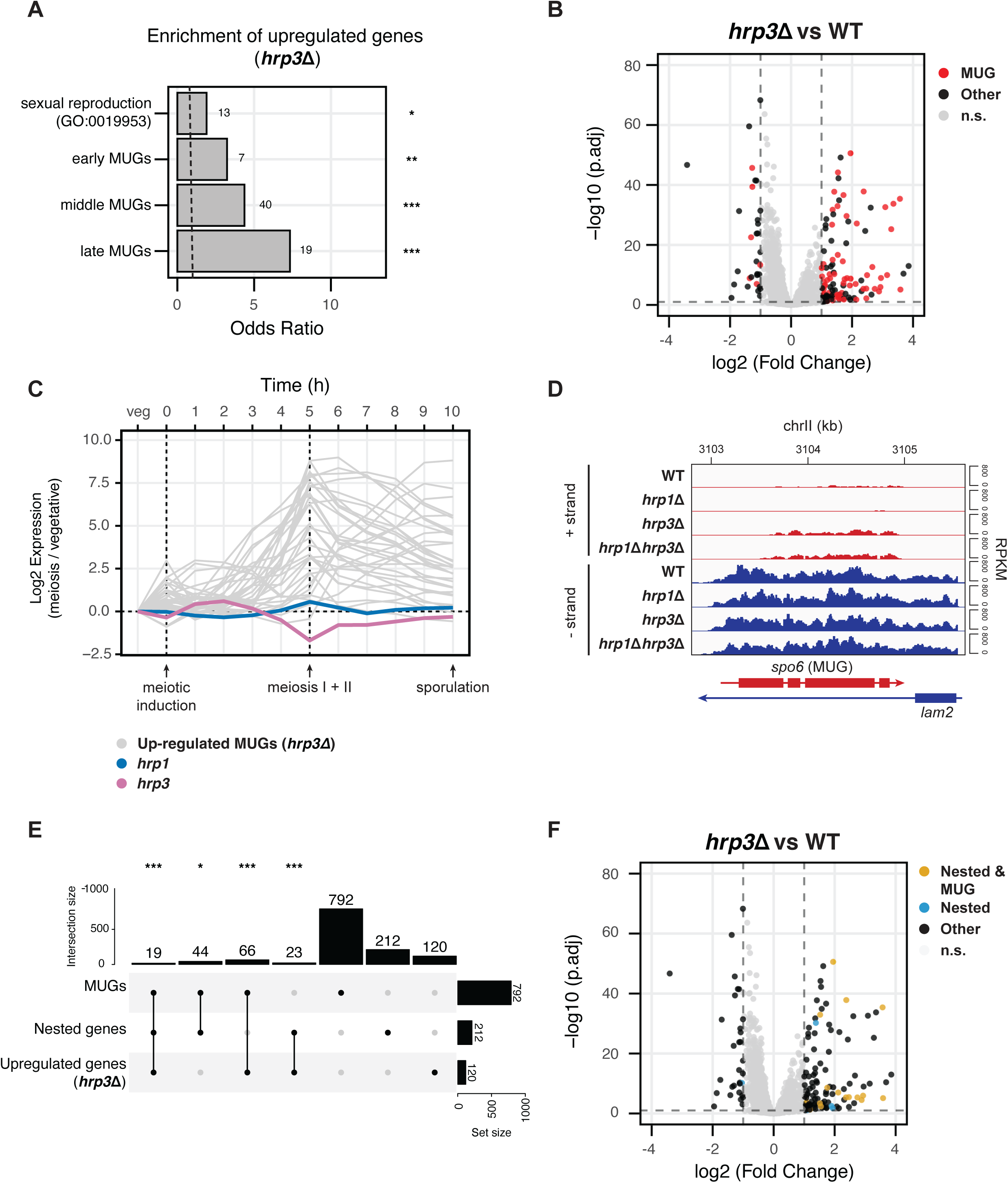
Hrp3-Mediated Antisense Transcription Regulates Meiotic and Nested Gene Expression in *S. pombe*. (A) Enrichment analysis of upregulated genes in the *hrp3*Δ mutant for the sexual reproduction GO term (GO:0019953) and early, middle, and late meiotically upregulated genes (MUGs) as reported by Mata et al. The dashed line indicates an odds ratio of 1. Statistical analysis was performed using Fisher’s exact test. Asterisks indicate statistical significance: p < 0.05 (*), p < 0.01 (**), p < 0.001 (***), n.s. (not significant). (B) Volcano plot representing the differentially expressed genes in *hrp3*Δ versus WT. MUGs are highlighted in red, while all other genes are in black. Genes in grey do not meet the thresholds for both significance (-log_10_(p.adj) > 1) and effect size (−1 < log_2_FC < 1). (C) Line plot showing the temporal expression of *hrp1*, *hrp3*, and 35 MUGs upregulated in *hrp3*Δ at 1-hour intervals during meiosis in *S. pombe*. Data were obtained from Mata et al.^71^ (D) Genome browser track of mRNA-seq data for the *spo6* and *lam2* genes in WT, *hrp1*Δ, *hrp3*Δ and *hrp1*Δ*hrp3*Δ. Tracks are separated into the + strand (red) and – strand (blue), with values shown in RPKM. The MUG within the pair is denoted in brackets. (E) UpSet plot showing the overlap between MUGs, convergent nested genes and upregulated genes in *hrp3*Δ. Statistical analysis was performed using the hypergeometric test. Asterisks indicate statistical significance: p < 0.05 (*), p < 0.01 (**), p < 0.001 (***), n.s. (not significant). (F) Volcano plot representing the differentially expressed genes in *hrp3*Δ versus WT. Genes which are both convergently nested and MUGs (n = 19) are highlighted in gold, other convergent nested genes (n=193) are highlighted in blue, and all other genes are in black. Genes in grey do not meet the thresholds for both significance (-log_10_(p.adj) > 1) and effect size (−1 < log_2_FC < 1).

To test this, we established a phenotypic assay to assess the impact of widespread antisense transcription on cellular fitness. We assayed the growth of WT, *hrp1*11, *hrp3*11, and *hrp1*11*hrp3*11 under stress conditions targeting DNA damage repair (bleomycin^72^), cell cycle/chromatin (caffeine^73^), the anti-fungal agent clotrimazole^74^, protein folding (guanidinium hydrochloride^75^), and transcription and replication (mycophenolic acid^76^) (**Fig. 5A; Group I**). Our results revealed that *hrp3*11 and *hrp1*11*hrp3*11 exhibited significantly reduced growth under all tested stress conditions compared to WT, with particularly pronounced sensitivity to caffeine, bleomycin, and mycophenolic acid. In contrast, *hrp1*11 exhibited minimal to no growth defects relative to WT. This pattern mirrors the extent of nucleosome disruption and antisense transcription in these mutants, suggesting that elevated antisense transcription imposes a broad fitness cost.

**Figure 5.**
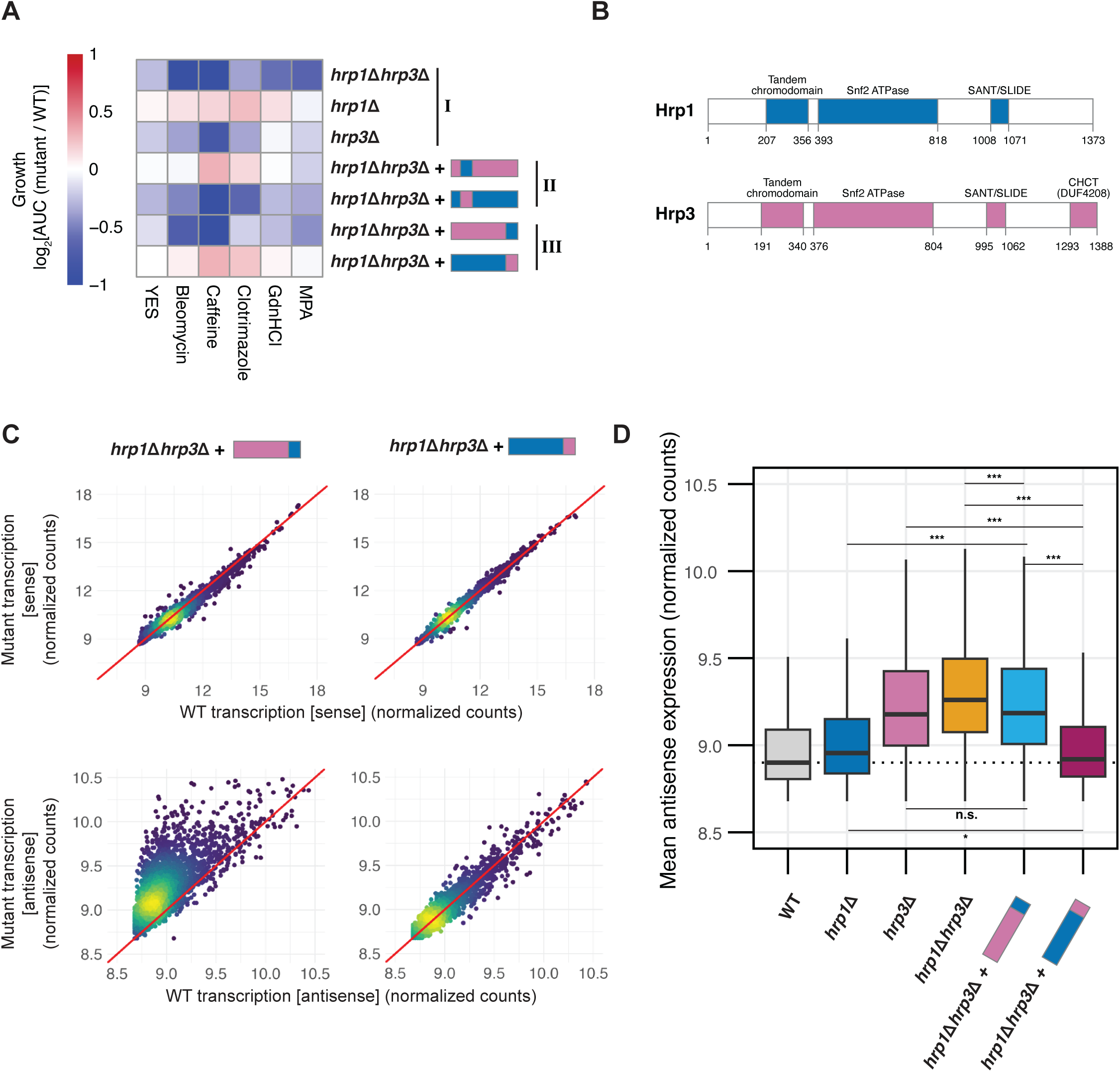
The CHCT Domain of Hrp3 Prevents Antisense Transcription. (A) Heatmap of growth phenotypes across four biological replicates when grown in YES medium alone or supplemented with 25 ng/μl bleomycin, 14 mM caffeine, 200 ng/ml clotrimazole, 5 mM guanidine hydrochloride (GHCl), or 25 μg/ml mycophenolic acid (MPA), represented by log_2_AUC values of mutants relative to WT. Roman numerals indicate mutant groups: (I) deletion mutants (II) chromodomain-swapped chimeras in the *hrp1Δhrp3Δ* mutant background, and (III) C-terminus-swapped chimeras in the *hrp1Δhrp3Δ* mutant background. Data represent the mean growth from four biological replicates. (B) Schematic representation of the protein structures of Hrp1 and Hrp3, including domain annotations with residue start and end positions. (C) Density-colored scatterplots representing mutant (*hrp1Δhrp3Δ::hrp3-CT_hrp1_ or hrp1Δhrp3Δ::hrp1-CT_hrp3_*) transcription against WT transcription for either sense transcripts (top row) or antisense transcripts (bottom row) for all protein-coding genes. Count data is derived from variance-stabilizing transformation (vst) of raw mRNA-seq counts from three biological replicates. (D) Boxplot of normalized antisense expression counts (as calculated in C) for WT, *hrp1Δ, hrp3Δ*, *hrp1Δhrp3Δ, hrp1Δhrp3Δ::hrp3-CT_hrp1_ and hrp1Δhrp3Δ::hrp1-CT_hrp3_*. The dotted line represents the median antisense expression in WT. Statistical analysis was performed using ANOVA with Tukey’s Honestly Significant Difference (HSD) test. Asterisks indicate statistical significance: p < 0.05 (*), p < 0.01 (**), p < 0.001 (***), n.s. (not significant).

### The CHCT Domain of Hrp3 Prevents Antisense Transcription

We next investigated the molecular basis for Hrp3’s suppression of antisense transcription. To do this, we leveraged the functional contrast with Hrp1, which exhibited a dramatically smaller antisense phenotype (**Fig. 1D**). We first compared their protein sequences and found the greatest divergence in their C-terminal regions (**Fig. S6A**). Detailed annotation using a Pfam search identified the CHD1 helical C-terminal (CHCT) domain^77^ as a domain of unknown function (DUF4208; residues 1293–1388) in Hrp3, which was absent in Hrp1 (**Fig. 5B**). Structural predictions using AlphaFold showed that the CHCT domain of Hrp3 forms a structured bundle of five α-helices (**Fig. S6B, S6D**), consistent with previous NMR spectroscopy studies^77^, and is highly conserved among CHD1-family proteins in other eukaryotes. In contrast, the C-terminus of Hrp1 had no predicted structure (**Fig. S6C, S6E**) and has diverged from Hrp3 within the fungal *Schizosaccharomyces* lineage^78^. These findings highlight the structural and evolutionary divergence of the C-terminal regions, which likely underlies the functional differences between Hrp1 and Hrp3.

To test whether the CHCT domain is a key driver of Hrp3’s impact on antisense transcription, we utilized the phenotypic assay described earlier and generated four chimeric mutants by swapping either the chromodomains (CD) or C-termini (CT) between the two paralogs, knocking them into the *hrp1*11*hrp3*11 background (**Fig. S6F**). In this assay, if Hrp3 function is complemented, then chimeras should resemble the *hrp1*11, whereas if they do not, then they should resemble the *hrp3*11. Chromodomains were chosen as a control because, unlike the C-termini, they are relatively conserved between Hrp1 and Hrp3 (**Fig. S6A**). Growth assays under stress conditions revealed that swapping the chromodomains between Hrp1 and Hrp3 did not alter the phenotype, indicating functional interchangeability (**Fig. 5A; Group II**). By contrast, replacing the Hrp3 C-terminus with that of Hrp1 (*hrp1*11*hrp3*11*::hrp3-CT_hrp1_*) recapitulated the growth defects of *hrp3*11 (**Fig. 5A; Group III**), while introducing the Hrp3 CHCT domain onto Hrp1 (*hrp1*11 *hrp3*11*::hrp1-CT_hrp3_*) was sufficient to rescue the growth phenotype, even in the absence of Hrp3 (**Fig. 5A; Group III**), suggesting that the CHCT domain plays a critical role in Hrp3 function.

To determine whether the CHCT domain is involved in regulating antisense transcription, we next performed mRNA-seq on the *hrp1*11*hrp3*11*::hrp3-CT_hrp1_* and *hrp1*11 *hrp3*11*::hrp1-CT_hrp3_* mutants (**Fig. 5C, 5D**). Consistent with the growth phenotypes, transplanting the C-terminus of Hrp1 onto Hrp3 (*hrp1*11*hrp3*11*::hrp3-CT_hrp1_*) resulted in elevated levels of antisense transcription, closely resembling the *hrp3Δ* mutant. By contrast, swapping the CHCT domain of Hrp3 onto Hrp1 (*hrp1*11*hrp3*11*::hrp1-CT_hrp3_*) did not lead to such an increase. These findings indicate that the CHCT domain of Hrp3 is essential for the regulation of antisense transcription in cells.

### The CHCT Domain of Hrp3 Interacts with the Paf1 Complex via Prf1

Although not fully resolved in cryo-electron microscopy studies^79,80^, the terminal location of the CHCT domain of CHD1, distal from the nucleosome, suggested it may mediate protein-protein interactions. To investigate this possibility, we performed an *in silico* interaction screen using Alphafold2 Multimer^81^ on the CHCT domain with a set of 2,692 nuclear genes in *S. pombe*^82^. Our results identified 12 candidates with an average interface predicted template modelling (IPTM) score of > 0.5 (**Fig. S7A, Table S2**). We filtered the predicted interactors to include only proteins associated with transcription elongation, as CHD1 is primarily involved in this process. Among these, the RNAPII-associated Paf1 complex subunit Prf1 (RTF1 in other eukaryotes) emerged as the strongest candidate (ipTM = 0.5516). The same interaction was predicted between full-length Prf1 and Hrp3 (**Fig. S7B, S7C**), consistent with previous reports identifying the Paf1 complex as a potential CHD1 interactor in budding yeast^83,84^. In contrast, a similar prediction for Prf1 and Hrp1 did not reproduce the interaction (**Fig. S7D**). The predicted Hrp3–Prf1 interface consisted of a distal binding region and a hydrophobic pocket within the CHCT domain, comprising three contact points (R2, R3, R4) and a distal region (R1) that surround a small hydrophobic α-helix at the N-terminus of Prf1 (N-helix) (**Fig. 6A, 6B**). These contact points are conserved across major eukaryotic lineages (**Fig. S8A, S8B**). Further AlphaFold predictions with CHD1 from *Homo sapiens* and *Arabidopsis thaliana* also supported an interaction between the CHCT domain and the RTF1 N-terminal helix (**Fig. S7E–H**), suggesting possible evolutionary conservation of this mechanism.

**Figure 6.**
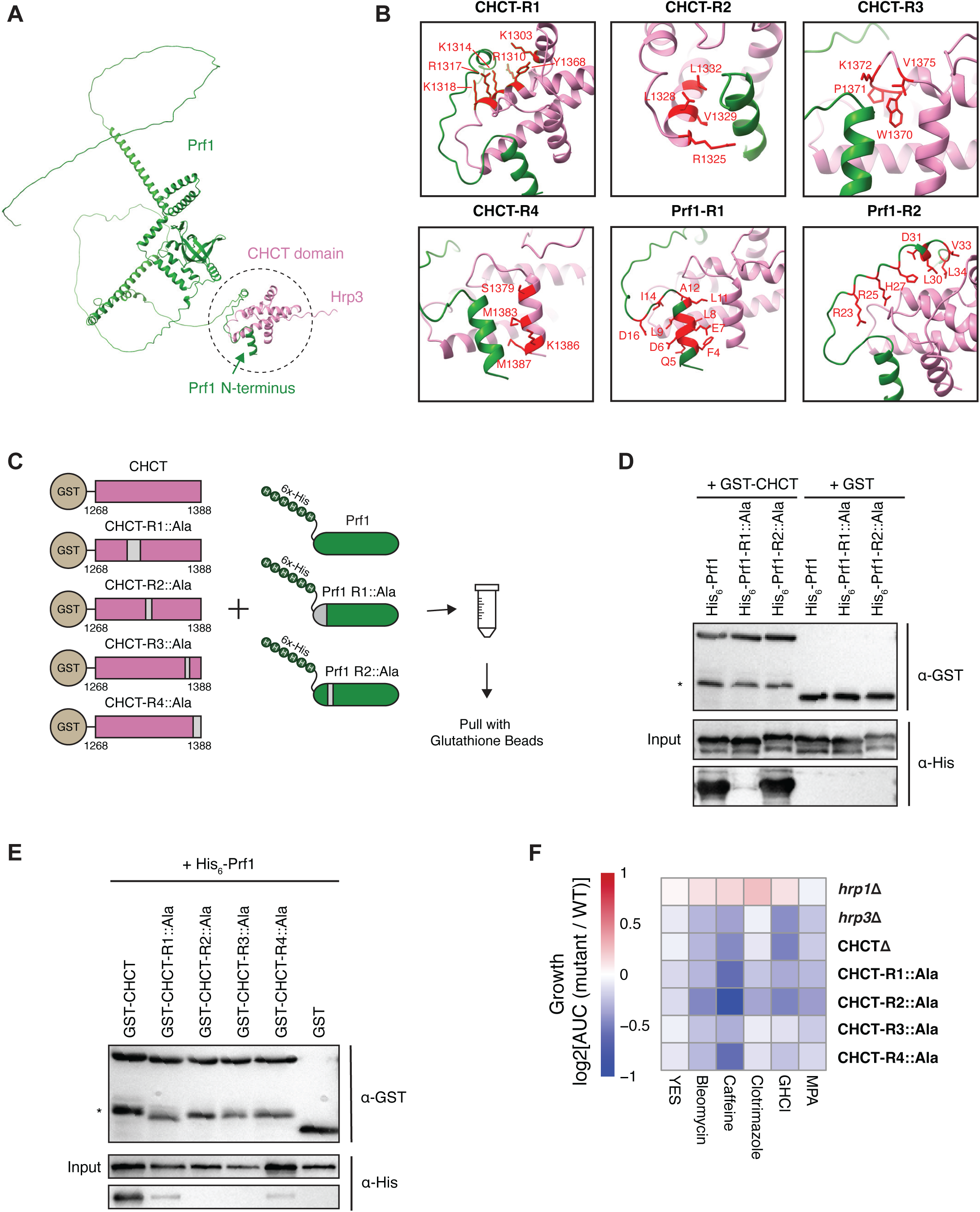
The CHCT Domain of Hrp3 Interacts with the Paf1 Complex via Prf1. (A) AlphaFold2-predicted structure of the interaction between the N-terminus of Prf1 and the CHCT domain of Hrp3. The predicted binding interface is highlighted with a dotted circle. (B) Close-up views across six AlphaFold2-predicted binding regions between Prf1 (green) and the CHCT domain (pink). Residues involved in the interaction are highlighted in red for each region. (C) Schematic of the in vitro interaction assay. Binding was tested between GST-tagged CHCT (wild-type and alanine mutants) and His-tagged Prf1 (wild-type and alanine mutants). Regions of the proteins with residues mutated to alanine are shown in grey. (D) Western blot showing anti-GST and anti-His signals for GST-pulldowns of GST-CHCT (wild-type or mutated) and wild-type His-Prf1. Asterisk (*) denotes truncated GST-CHCT constructs. (E) Western blot showing anti-GST and anti-His signals for GST-pulldowns for wild-type GST-CHCT with His-Prf1 (wild-type or mutated). Asterisk (*) denotes truncated GST-CHCT constructs. (F) Heatmap of growth phenotypes of *hrp1*Δ*, hrp3*Δ*, hrp3-CHCT*Δ, and *in vivo* alanine-swapped CHCT mutants (as in panel C) when grown in YES medium alone or supplemented with 25 ng/μl bleomycin, 14 mM caffeine, 200 ng/ml clotrimazole, 5 mM guanidine hydrochloride (GHCl), or 25 μg/ml mycophenolic acid (MPA). Data represent the log_2_AUC values of mutant growth relative to WT under various stress conditions across four biological replicates.

To experimentally test the Hrp3-Prf1 interaction predicted by the model, we recombinantly expressed GST-tagged CHCT along with both wild-type and mutated versions of His-tagged Prf1. In the mutated versions, residues within either the N-terminal helix or the distal region identified from AlphaFold analyses were substituted with alanine (**Fig. 6C**). Using an *in vitro* pulldown assay with the GST tag (and GST-only as a negative control), we found that His-Prf1 was pulled down by the CHCT domain, confirming their interaction. However, mutating the interacting residues in the Prf1 N-terminal helix to alanine, but not those in the distal region, abolished this interaction (**Fig. 6D**). Conversely, when we tested wild-type Prf1 against mutated versions of the CHCT domain (with alanine substitutions in the R1, R2, R3, and R4 regions), we observed a loss of binding to Prf1 (**Fig. 6C, 6E**). Further, when we extended these *in vitro* assays to test the CHCT mutants’ functional impact *in vivo*, strains expressing Hrp3-CHCT domain mutants displayed stress sensitivity comparable to *hrp3*11 and CHCT-deleted strains (**Fig. 6F**). These results demonstrate that the CHCT domain of Hrp3 is essential for its interaction with Prf1 and is critical for maintaining cellular fitness under stress conditions.

### Hrp3 and Prf1 Synergize to Regulate H2Bub Deposition, Nucleosome Organization, and Antisense Transcription

Given that RTF1 is enriched across gene bodies and is essential for H2Bub deposition via its histone modifying domain (HMD)^85^, we hypothesized that the Hrp3–Prf1 interaction regulates H2Bub deposition specifically during transcription elongation. Supporting this, Hrp3 localized across gene bodies with enrichment profiles similar to the profiles of the elongation marks H3K36me3 and H2Bub (**Fig. 7A**), and this localization was correlated with gene expression (**Fig. 7B**). In contrast, while Hrp1 also exhibited transcription-dependent enrichment (**Fig. S9A**), it was primarily localized at transcription termination sites (TTS) rather than gene bodies (**Fig. 7A, Fig. S9A**). Correlation analysis revealed a significant positive relationship between Hrp1 and Hrp3 enrichment over genes (**Fig. S9B**), suggesting that while the localization of both remodelers is associated with transcription, the CHCT domain of Hrp3 is critical for its association with transcription elongation along the gene body, whereas Hrp1, which lacks the CHCT domain, has diverged to perform a distinct function at the TTS. Notably, in the absence of Hrp3, Hrp1 may partially compensate for the suppression of antisense transcription, explaining the mild impact of the *hrp1*Δ mutant and the stronger impact of the *hrp1*Δ*hrp3*Δ double mutant compared to *hrp3*Δ alone on antisense production (**Fig. 3F**). However, this minor compensatory role of Hrp1 does not depend on factors associated with the CHCT domain, which is unique to Hrp3. Consistently, *hrp3*Δ and *hrp1*Δ*hrp3*Δ mutants exhibited a significant loss of H2Bub across gene bodies, while *hrp1*Δ mutants did not (**Fig. 7C**). Highly expressed genes also exhibited slightly higher loss of H2Bub (**Fig. S9C**), further supporting that Hrp3- and Prf1-mediated H2Bub deposition is tightly linked to transcription elongation.

**Figure 7.**
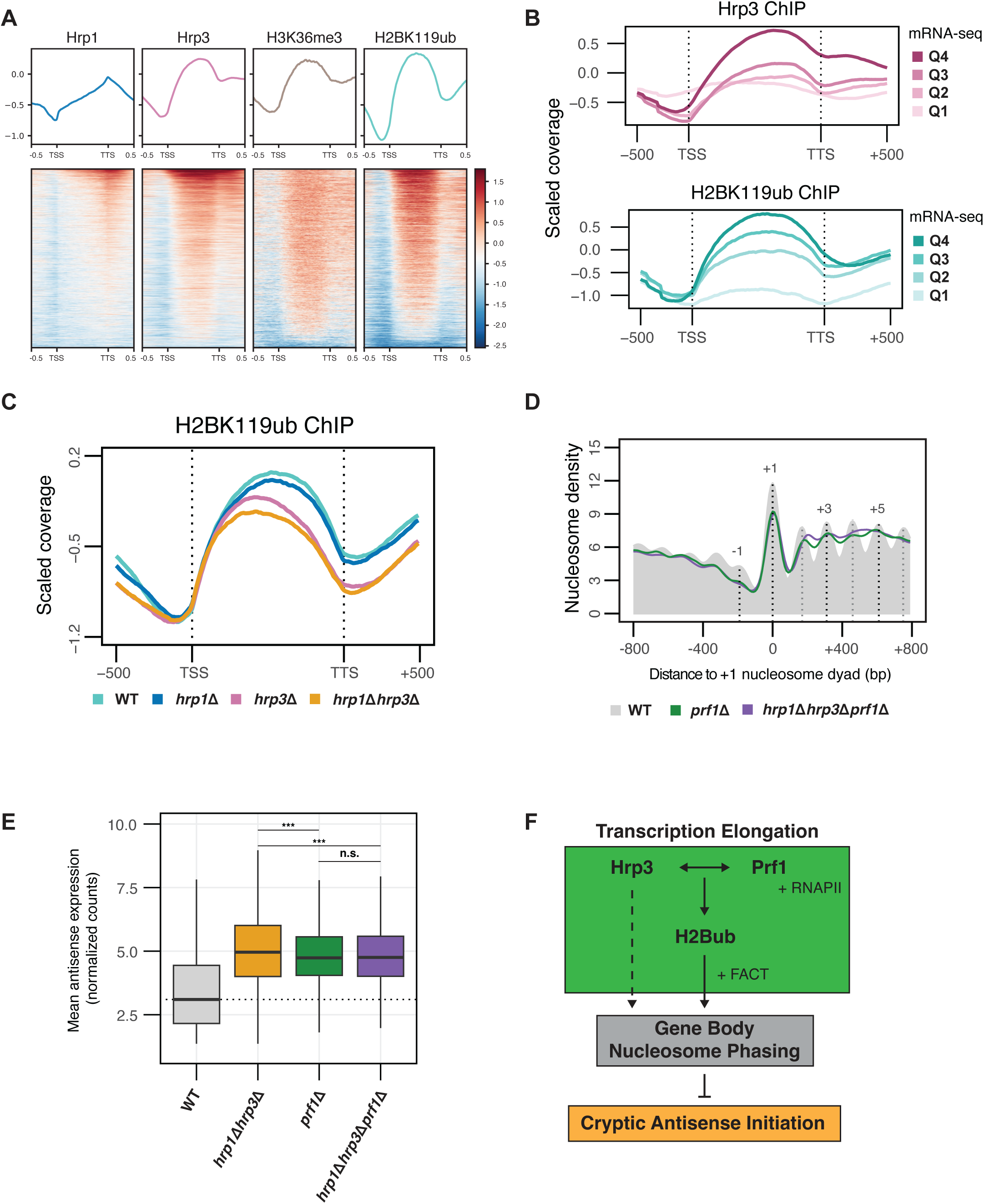
Hrp3 and Prf1 Synergize to Regulate H2Bub Deposition, Nucleosome Organization, and Antisense Transcription. (A) Metagene profiles and heatmaps showing ChIP-seq enrichment over input for Hrp1, Hrp3, H3K36me3 and H2BK119ub across all protein-coding genes. For Hrp1 and Hrp3, data were obtained using an anti-myc antibody in WT strains carrying myc-tagged Hrp1 and Hrp3. (B) Metagene plot of the ChIP-seq coverage relative to input for Hrp3-myc and H2BK119ub in WT across all protein coding genes. Genes were divided into four quartiles based on expression in WT. Quartiles are numbered by expression level, with Q1 being the lowest and Q4 being the highest. (C) Metagene plot of H2BK119ub ChIP-seq coverage relative to input for WT, *hrp1*Δ*, hrp3*Δ and *hrp1*Δ*hrp3*Δ across all protein coding genes. (D) Metaplots of normalized nucleosome density from MNase-seq centered on the +1 nucleosome dyad for WT, *prf1*Δ and *hrp1*Δ*hrp3*Δ*prf1*Δ. The WT nucleosome profile is shown in solid grey, with dotted lines indicating the positions of select nucleosome maxima (−1, +1, +3 and +5). Profiles include 800 bp upstream and downstream of the +1 nucleosome dyad for all protein-coding genes. The plotted data represents the average signal from two biological replicates. (E) Boxplot of normalized antisense expression counts for all antisense genes (calculated using vst-normalized raw mRNA-seq counts) for WT, *hrp1*Δ*hrp3*Δ*, prf1*Δ and *hrp1*Δ*hrp3*Δ*prf1*Δ. The dotted line represents the median antisense expression in WT. Data are calculated from three biological replicates. Statistical analysis was performed using ANOVA with Tukey’s Honestly Significant Difference (HSD) test. Asterisks indicate statistical significance: p < 0.05 (*), p < 0.01 (**), p < 0.001 (***), n.s. (not significant). (F) Schematic illustrating the relationship between Hrp3 and Prf1 in the context of H2Bub deposition and nucleosome phasing, and its impact on antisense transcription.

As H2Bub is known to enhance nucleosome stability^86^, we next assessed the impact of the loss of *prf1* on nucleosome organization using MNase-seq. Nucleosome phasing was severely disrupted in the *prf1*Δ mutant, as well as in the *hrp1*Δ*hrp3*Δ*prf1*Δ triple mutant (**Fig. 7D**). Given the similar degree of nucleosome disruption observed in both mutants, this suggests that Hrp3 and Prf1 are epistatic and function within the same pathway to regulate nucleosome spacing within gene bodies. Additionally, both the *prf1*Δ and *hrp1*Δ*hrp3*Δ*prf1*Δ mutants exhibited increased antisense transcription, comparable to the *hrp1*Δ*hrp3*Δ mutant (**Fig. 7E**), further supporting the idea that Hrp3 and Prf1 act together in the same pathway to control antisense transcription (**Fig. 7F**). Consistent with the observations upon loss of *hrp3*, sense transcription was also largely unaffected in *prf1*Δ and *hrp1*Δ*hrp3*Δ*prf1*Δ (**Fig. S9D, S9E**). However, the *prf1*Δ and *hrp1*Δ*hrp3*Δ*prf1*Δ mutants displayed much stronger growth defects than the *hrp1*Δ*hrp3*Δ mutant, indicating that Prf1 has additional essential roles in cells^87^ (**Fig. S9F**). Together, these findings highlight the cooperation between Hrp3 and Prf1 in establishing nucleosome organization, which is critical for maintaining proper gene regulation in cells.

## DISCUSSION

In this study, we demonstrate that the CHD1-family nucleosome remodeler Hrp3 is a central regulator of transcription in *Schizosaccharomyces pombe*. Hrp3 establishes nucleosome arrays within gene bodies, counteracting DNA sequence biases that promote cryptic antisense initiation by exposing embedded promoter-like regions. These antisense transcripts include a subset of nested meiotic genes that leverage Hrp3’s regulatory function to suppress their expression during vegetative growth. The establishment of regular nucleosomes arrays over gene bodies is mediated by the combined action of Hrp3’s Snf2-ATPase and CHCT domains. The CHCT domain facilitates interaction with Prf1, a subunit of the Paf1 elongation complex, effectively linking the elongation machinery to the SWI/SNF nucleosome remodelers. This interaction enables precise spatial and temporal control of nucleosome positioning, ensuring proper chromatin organization and transcriptional regulation. Together, these findings underscore the role of Hrp3 in maintaining chromatin organization and proper gene expression, while also highlighting the coordination of regulatory mechanisms spanning chromatin structure and gene expression required for proper promoter licensing in cells.

Antisense transcription is an evolutionarily ancient and ubiquitous phenomenon, observed throughout life^5,88^, but its evolutionary significance and regulatory mechanisms remain incompletely understood. Furthermore, the evolutionary pressures governing antisense transcription remain a subject of debate^12,89^, as its extent appears to be highly species-specific. For instance, 30–40% of genes in humans are bidirectionally transcribed, while this proportion increases to up to 70% in mice^8,90–92^. Here, we expand the catalog of antisense transcripts in *S. pombe*, identifying thousands of dormant antisense RNAs that overlap up to 55% of all protein coding genes. Many of these antisense transcripts, including several meiotic genes, are nested within larger host genes in a convergent orientation and repressed by Hrp3. While most of these antisense transcripts do not appear to directly regulate the sense expression of overlapping genes, as observed in other species^10,12,93,94^, they may serve alternative *trans*-regulatory roles or encode functional proteins. This suggests that nature may have co-opted these additional transcripts for diverse functions and regulatory roles in a species-specific manner, akin to how gene duplication and neofunctionalization have driven evolutionary innovation^95–98^. In this way, antisense transcription can be viewed as a source of complexity, providing raw material for the evolution of novel regulatory networks and functional elements. The dual nature of antisense transcription—acting as either beneficial regulatory elements or harmful transcriptional noise—underscores the complexity of transcriptional regulation and the evolutionary pressures that maintain this phenomenon. Unregulated antisense transcription, as seen in *hrp3*Δ mutants, imposes significant fitness costs, particularly under stress conditions, emphasizing the necessity of tight regulatory control.

A key question arising from our study is how Hrp3 achieves the precise regulation of nucleosome positioning in coordination with the transcriptional machinery. Our study, together with recent work in budding yeast^99^, identifies the interaction between the Hrp3 CHCT domain and the elongation complex subunit Prf1 as a critical link in this process. Disruption of this interaction—either by loss of Hrp3, the CHCT domain, or Prf1—results in similar phenotypes, including impaired nucleosome phasing and increased antisense transcription. These findings align with previous studies showing that H2Bub, deposited by the Paf1 complex, facilitates the activity of FACT^86,100^, a histone chaperone that reassembles nucleosomes in the wake of RNAPII. Loss of FACT similarly disrupts nucleosome spacing and increases antisense expression^101–105^, supporting a model in which Hrp3, Prf1, and FACT act together during transcription elongation to ensure proper nucleosome spacing and the suppression of cryptic transcription, underscoring the intricate crosstalk required between nucleosome remodelers and the transcription machinery. This also helps explain why antisense transcription is overwhelmingly enriched to initiate within gene bodies, as it provides a direct connection between the spatial context of transcription elongation, nucleosome positioning, and the regulation of antisense initiation.

Additionally, our findings show that chromatin, specifically well-phased nucleosome arrays, play a key role in regulating antisense transcription. Building on previous studies that established the role of nucleosome arrays in suppressing cryptic transcription^32–34^, we extend this concept to within gene bodies. We show that cryptic promoters can arise within canonical protein coding genes, which Hrp3 works together with the transcription apparatus to suppress.

The evolution of nucleosome arrays in eukaryotes, alongside the diversification of nucleosome remodeler families as their architects, likely reflect the demand for the increasing complexity of transcriptional regulation during eukaryogenesis^106^. Furthermore, the widespread distribution of AT-rich regions which can function as cryptic antisense promoters within genes appears to be an inevitable consequence of evolutionary constraints, such as the local nucleotide composition imposed by protein-coding sequences. Indeed, studies have shown that up to 36% of transcription factor binding sites in humans lie within genes^107^. Although codon redundancy may have partially evolved to mitigate this issue^108,109^, the nucleosome remodeler Hrp3 remains essential for ensuring that these cryptic promoters remain latent unless specifically required. This also suggests that the widespread distribution of cryptic antisense promoters is an inherent feature of eukaryotic genomes, shaped by both sequence composition and chromatin architecture.

In conclusion, our study highlights the critical role of Hrp3 in maintaining proper promoter licensing and antisense transcription regulation in *S. pombe*. By positioning nucleosomes to occlude cryptic antisense promoters and coordinating with the transcription elongation machinery, Hrp3 ensures proper gene regulation and cellular fitness. The thousands of newly identified antisense transcripts revealed in this study likely play context-specific roles that remain to be fully characterized. These findings offer valuable insights into the evolutionary and functional importance of nucleosome remodelers and their pivotal role in shaping the transcriptional landscape of eukaryotic genomes.

## RESOURCE AVAILABILITY

Further information and requests for resources and reagents should be directed to and will be fulfilled by the lead contact, Frédéric Berger.

### Data and code availability

Datasets from the analyses are provided as supplementary data files. The annotated list of antisense transcripts identified in this study are available as a GFF3 file at https://github.com/Gregor-Mendel-Institute/kok_2025/tree/main/output/annos. NGS data are deposited in GEO under accession numbers: PRO-seq (GSE302374), MNase-seq (GSE302388), mRNA-seq (GSE302386), and ChIP-seq (GSE302387). All other raw and processed data are available upon request to the lead contact.

## ACKNOWLEDGEMENTS

We thank the entire Berger group, for their insightful, considerate, and helpful discussions, as well as Matt Watson for editorial assistance during the preparation of the manuscript. We are especially grateful to Hwan Su Yoon (Sungkyunkwan University, Korea) for their support with long-read transcriptomics and to Dominik Handler for providing scripts for AlphaFold. We also acknowledge the Molecular Biology Service and Media Kitchen for their constant supply of plates, basic reagents, and cloning and sequencing services. Additionally, we thank the Vienna BioCenter Core Facilities, in particular the Next Generation Sequencing facility for their advice and swift handling of all our requests and helpful discussions. For open access purposes, the authors have applied a CC BY public copyright license to any author accepted manuscript version arising from this submission.

## AUTHOR CONTRIBUTIONS

Conceptualization: FB and JYK; Methodology: JYK and ZHH; Validation: JYK and EA; Formal Analysis: JYK and EA; Investigation: JYK, EA, and FB; Data Curation: JYK and EA; Writing – original draft: JYK and FB; Supervision: ZHH and FB; Project Administration: FB; Funding Acquisition: ZHH and FB.

## SUPPLEMENTAL INFORMATION

Supplementary Data

**Figures S1-S9.**

**Table S1.** Early, middle and late MUGs upregulated in *hrp3Δ*.

**Table S2.** Top candidates for predicted interaction with the CHCT domain of Hrp3.

**Table S3.** *Schizosaccharomyces pombe* strains used in this study.

## MATERIALS AND METHODS

### Yeast Culture, Strain Manipulation, Phenotypic and Mating Assays

All strains used in this study are listed in **Table S3**. Yeast (*S. pombe* strain 972) were cultured under standard conditions (32 °C). Unless otherwise stated, media used were either EMM2 (Edinburgh Minimal Media)^110^ or YES (Yeast Extract with Supplements)^111^.

*S. pombe strains* were manipulated either using one-step PCR-based HR mutagenesis^112^ or the SpEDIT CRISPR editing platform^113^. For both, heat-shock was used to transform chemically-competent parent strains^114^. Donor templates for homologous recombination were prepared using either Gibson assembly^115^ using reagents supplied by the IMP Molecular Biology Service combined, with site directed mutagenesis, or by direct synthesis from commercial vendors (IDT, Genscript). For commercially synthesized HR templates, sequences were codon optimized according to *S. pombe* codon usage using Benchling (https://benchling.com). In all experiments, WT control transformations were performed such that synonymous mutations were introduced to delete CRISPR gRNA target sites. All manipulations were verified by PCR genotyping and where appropriate sequencing of the relevant locus.

To construct the *ura4*-based reporter, a construct (PJY_135) consisting of homology to the 5**′** of the *ura4*-D18 locus^116^, the selectable marker natMX6, a ∼1 kb fragment corresponding to either the S. pombe *adh1* promoter, a ∼500 bp fragment tested to have minimal transcriptional activity, or ∼500 bp fragments corresponding to antisense promoters within *rap1*, *atg9*, *crt10*, *spb70* or *orc4* were amplified from gDNA extracted from WT *S. pombe*, and the *ura4*^+^ gene were assembled by Gibson. The *ura4-D18* locus was then replaced with the reporter via one-step PCR followed by homologous recombination^112^. Correct integration was verified by PCR genotyping.

Phenotypic assays in this study were conducted using a plate-based bulk growth assay. OD_600_ was continuously monitored in 384-well plates (Nunc) under the stress conditions specified in the figure legends, at the standard growth temperature with shaking. Saturated cultures were diluted 100-fold for sub-culturing, and all assays were performed in either YES, EMM complete, or EMM without uracil (EMM-Ura, for reporter assays). Data collection was carried out using a BioTek Epoch2 microplate reader. Growth curves were analyzed in R, and the area under the curve (AUC) was calculated using the Growthcurver package (https://cran.r-project.org/web/packages/growthcurver/vignettes/Growthcurver-vignette.html). All subsequent data analysis was done in R. For spotting assays on agar plates, saturated cultures were diluted 1:10, and their OD_600_ was adjusted to 0.6. Subsequently, 1:2 serial dilutions were prepared, and cells were spotted onto YES agar plates using a 48-prong spotter.

### Bacterial protein expression and in vitro pulldown experiments

Sequences corresponding to WT and mutated versions of *prf1* and the *hrp3* CHCT domain were synthesized directly from commercial vendors (Genscript) and amplified by PCR. The *prf1* sequences were cloned into pET28a (Novagen), while the *hrp3* CHCT domain sequences were cloned into pGEX-4T-1 (Cytiva). Assembly of all plasmids was carried out by Gibson.

The plasmids expressing WT and mutated versions of 6xHis-tagged *prf1* were then co-transformed with plasmids expressing WT or mutated versions of GST-tagged *hrp3* CHCT domain into E. coli BL21 (DE3) RIL. Ten milliliter cultures grown overnight at 37 °C were diluted into 200 ml of fresh LB medium and grown at RT for 3 h and then induced for 5 h at RT. Cells equivalent to 100 of culture were resuspended in 5 ml of TBS containing 0.1% Triton X-100, 1 mM DTT, protease inhibitors (Roche), 10 µl of benzonase (2 µg/ml) and lysozyme (50 mg per 50 ml). After sonication for 10 min at high intensity (10”on/15”off) and 5 min at medium intensity (10”on/15”off) extracts were centrifuged for 20 min at 4 °C at 40,000×g. Extracts were incubated with 50 µl of magnetic Glutathione (Thermo Fisher Scientific) or Protein A (Cytiva) beads (each with 2.5 ml of protein extract) at RT for 2 h. Beads were collected and washed six times with extraction buffer (without benzonase and lysozyme) and finally resuspended in 80 µl of 1× SDS-PAGE loading buffer prepared in extraction buffer and boiled for 5 min. Proteins were then separated on an 12% SDS-PAGE gel and analyzed by either colloidal blue staining or Western blot. For SDS-PAGE, separation was performed using self-cast gels using standard methods. Gels were stained using colloidal blue reagents (MBS Blue) provided by the Molecular Biology Service. For Western blots, proteins were transferred using a standard wet protocol to a 0.2 µm nitrocellulose membrane (Cytiva Cat. No. 10600004), blocking in 5% non-fat dry milk (Maresi) dissolved in PBST. Anti-His (Sigma-Aldrich, H1029) and anti-GST (Santa Cruz Biotechnology, sc-138) antibodies were used at 1:1000 diluted in blocking buffer and detected using chemiluminescence following incubation with anti-mouse HRP conjugates (Biorad Cat. No. 1706516), which were diluted 1:10,000 in blocking buffer. Imaging of gels and Western blots was performed using a Thermo Fisher iBright 1500 imaging system.

### mRNA Sequencing Library Preparation and Analysis

Total RNA was extracted from yeast strains grown to mid-exponential phase (OD_600_ ∼0.5) using a standard hot acid phenol protocol. Libraries were then prepared by polyA-tail enrichment using the NEBNext Ultra II Directional RNA Library Prep Kit for Illumina (NEB Cat. No. E7760L) with the polyA selection module (NEB Cat. No. E7490L). Size selection steps and final library cleanup were performed using the NA clean-up bead solution provided by the VBC core facilities, which is adapted from DeAngelis et al. 1995^117^ using carboxylate-modified Sera-Mag Speed beads (Cytiva). Library quality control was performed using the Advanced Analytical Fragment Analyzer using the HS NGS fragment analysis kit (Agilent Cat. No. DNF-474) as well as qPCR using reagents produced by the Molecular Biology Service in conjunction with commercial DNA standards (Roche Cat. No KK4903). Data was collected on a Thermo Scientific QuantStudio5 instrument.

Prepared libraries were sequenced on either an Illumina NextSeq2000 or NovaSeq S4 at the Vienna BioCenter Core Facilities Next Generation Sequencing Core using paired-end mode. Quality control of the data was performed using FastQC (Babraham Institute). Transcript-level quantification against the *S. pombe ASM294v2 reference genome* available from ENSEMBLv55 was performed using Kallisto^118^. Differential expression analysis was performed using DESeq2^119^ in R, where all subsequent data manipulation was performed.

### PRO-seq Library Generation and Analysis

We performed a variant of PRO-seq, qPRO-seq, as recently published^64^, with modifications^120^ to make it compatible with *S. pombe*. *S. pombe* cells were grown in 10 ml YES media to OD_600_ ∼0.4–0.5. Cells were then harvested by centrifugation at 400 x *g* for 5 min at 4 °C. Cells were then resuspended in 10 ml of ice-cold PBS, spun down once more, then resuspended in 10 ml ice-cold yeast permeabilization buffer (0.5% Sarkosyl, 0.5 mM DTT, Roche cOmplete protease inhibitor cocktail, 4U/ml Invitrogen RiboLock RNAse Inhibitor). After 20 min incubation on ice, cells were once again collected by gentle centrifugation, then resuspended in 50 µl storage buffer (10 mM Tris-HCl, pH 8.0, 25% (v/v) glycerol, 5 mM MgCl_2_, 0.1 mM EDTA, 5 mM DTT).

A 2X run-on reaction master mix (40 mM Tris pH 7.7, 64 mM MgCl2, 1 mM DTT, 400 mM KCl, 40 µM Biotin-11-CTP, 40 µM Biotin-11-UTP, 40 µM ATP, 40 µM GTP, 1% sarkosyl) was prepared and preheated to 30 °C. To each aliquot of permeabilized cells, 50 µl of master mix and 1 µl of RNAse inhibitor were added, then mixed by gently pipetting with wide-bore pipet tips before immediately incubating at 30 °C for 5 min. To ensure precise timing, samples were done in batches staggered by 30–60 sec. Using the Norgen RNA Extraction Kit (Cat. No. 37500), 350 µL of RL buffer were added to each sample and then vortexed. 240 µl of absolute ethanol were added, then the entire well-mixed solution was applied to the RNA extraction column. The column was spun at 3500 x *g* for 1 min at 25 °C, then washed once with 400 µl of wash solution A. This step was repeated a second time, then the column was dried by spinning at high speed for 2 min. RNA was eluted twice in 50 µl of DEPC-treated MonoQ water, the two fractions were pooled and then briefly denatured at 65 °C for 30 sec. After snap cooling on ice, 25 µl of ice cold 1 M NaOH were added to each sample, which was then incubated for 10 min on ice. Finally, samples were quenched with 125 µl of cold 1 M Tris-HCl pH 6.8, then cleaned up using a standard ethanol precipitation overnight. Samples were resuspended in a small volume of DEPC water (6 µl).

Meanwhile, 10 µl per sample of Dynabeads MyOne Streptavidin C1 beads (Thermo Fisher Cat. No. 65001) were prepared by washing once with bead preparation buffer (0.1 M NaOH, 50 mM NaCl), then once in bead binding buffer (10 mM Tris-HCl, pH 7.4, 300 mM NaCl, 0.1% (v/v) Triton X-100, 1 mM EDTA, RNAse inhibitors), then resuspended in 25 µl bead binding buffer.

3’ RNA adapter ligation was performed by adding 1 µl 10 µM oligo VRA3 to the resuspended RNA, boiling at 65 °C for 30 sec, then snap cooling on ice. Ligation reactions were then prepared with T4 RNA Ligase using the provided instructions. After incubating reactions at 25 °C for 1 hr, ligated RNA was bound to the 25 µl previously prepared streptavidin beads by adding 55 µl bead binding buffer and the entire ligation reaction. Samples were incubated for 20 min at 25 °C, then washed once with 500 µl high salt wash buffer (50 mM Tris-HCl, pH 7.4, 2 M NaCl, 0.5% (v/v) Triton X-100, 1 mM EDTA, 2 µl/10 ml Ribolock RNAse inhibitor), transferring beads to a clean tube. Beads were then washed once more in the new tube with low salt wash buffer (5 mM Tris-HCl, pH 7.4, 0.1% (v/v) Triton X-100, 1 mM EDTA, 2 µl/10 ml Ribolock RNAse inhibitor), then resuspended in 20 µl T4 PNK reaction mix (1X T4 PNK Buffer, 1 mM ATP, 1 µl T4 PNK, 1 µl RiboLock RNAse inhibitor). Samples were then incubate at 37 °C for 30 min, then the beads were collected by magnet and resuspended in 20 µl ThermoPol reaction mix (1X ThermoPol buffer, 1 µl RppH, 1 µl RiboLock RNAse inhibitor), then incubated at 37 °C for 1 hr. Beads were once again collected by magnet, then the 5’ RNA adaptor (REV5) was ligated using T4 RNA ligase, setting up the reaction using the provided manual and incubating at 25 °C for 1 hr. Beads were then washed once with high salt wash buffer, transferring to a clean tube, then once with low salt buffer, before being resuspended in 300 µl of TRIzol reagent (Ambion). Samples were vortexed for 20 sec, then incubated on ice for 3 min before adding 60 µl of chloroform, vortexing for 15 sec, then incubating on ice for a further 3 min. Samples were then spun at 20,000 x *g* for 8 min at 4 °C, then the upper ∼180 µl aqueous phase was transferred to a clean tube and precipitated using a standard ethanol precipitation.

Finally, pellets were resuspended in 13.5 µl reverse transcription resuspension mix (8.5 µl DEPC water, 4 µl 10 µM primer RP1, 1 µl 10 mM dNTPs), denaturing at 65 °C for 5 min, snap cooling on ice. Reverse transcription was then performed using Maxima H Minus RT (Thermo Fisher Cat. No. EP0753) by adding 6.5 µl RT master mix (4 µl 5X RT buffer, 1 µl DEPC water, 0.5 µl RiboLick RNAse inhibitor, 1 µl Maxima H Minus RT), then cycling as follows: 50 °C for 30 min, 65 °C for 15 min, 85 °C for 5 min. 2.5 µl of indexed RPI-n primer (10 µM) were then added to each RT reaction, then PCR was performed as a 100 µl reaction using the Q5 polymerase (NEB) with the included high GC content enhancer. Samples were then PCR-amplified to the beginning of the logarithmic phase, as determined using a small-scale qPCR reaction run in parallel (typically less than 20 cycles).

Libraries were sequenced as described previously at the Vienna BioCenter Core Facilities Next Generation Sequencing Facility. The data analysis was performed by adapting available pipelines^64^. Briefly, we created a Nextflow pipeline (https://github.com/Gregor-Mendel-Institute/PROalign) to perform all QC, alignment, and read processing steps, generating strand-specific bigwig files that were subsequently analyzed in R.

### Long-read transcriptomics (PacBio Iso-seq)

Total RNA was extracted from yeast strains grown to mid-exponential phase (OD600 ∼0.5) using a standard hot acid phenol protocol. For PacBio Iso-Seq, the sequencing libraries were prepared by DNA Link Sequencing Labs with the SMRTbell Barcoded Adapter Plate 3.0, followed by Iso-Seq sequencing on the Revio platform using SMRT Link v25.1. CCS reads (also known as HiFi reads) and high-quality isoforms were identified using the Iso-Seq4 (https://github.com/PacificBiosciences/IsoSeq) analytic software with a default parameter.

Subsequent analyses were performed in-house as follows: the clustered transcripts were mapped to the *Schizosaccharomyces pombe* genome using minimap2 v2.24^121^ with the parameters “-ax splice:hq-uf”. Redundant or highly similar transcripts were collapsed using TAMA Collapse with parameter “-a 100 –m 100 –x no_cap”. Collapsed transcript sets from the two independent libraries were subsequently merged using TAMA Merge^122^ with parameter “-d merge_dup”.

To identify antisense transcripts, we downloaded the GFF3 file containing gene annotations from the <u>Ensembl FTP</u> (https://ftp.ensemblgenomes.ebi.ac.uk/pub/fungi/release-60/gff3/schizosaccharomyces_pombe/). Features annotated as “antisense” and/or “non-coding” were excluded. The resulting reduced GFF3 file was then used as a reference for gffcompare^123^. Long-read transcripts classified by gffcompare as “exonic overlap on the opposite strand” (class code ‘x’) were directly adopted as *our* antisense transcript annotation.

### MNase-seq Library Generation and Analysis

We performed MNase-seq by adapting a published protocol^124^. *S. pombe* strains (100 ml) were grown to mid-exponential phase (OD_600_ ∼0.4-0.5) in YES, fixed with 0.5% formaldehyde for 20 min at RT, quenched with 125 mM glycine for 10 min, then collected by centrifugation. Cells were then washed with 10 ml dH_2_O, followed by resuspension into 5 ml of ice cold preincubation solution (20 mM citric acid, 20 mM Na_2_HPO_4_, 40 mM EDTA pH 8.0, 30 mM β-mercaptoethanol), and incubation in a 30 °C water bath for 10 min. Cells were next harvested and resuspended in 2 ml Sorbitol/Tris buffer (1 M sorbitol, 50 mM Tris/HCl pH 7.5, 10 mM β-mercaptoethanol), and 2 mg of Zymolyase 100T lyophilized powder (Nacalai Tesque) was resuspended in 40ul of water and added per 100 ml culture to produce spheroplasts. Samples were then incubated in a 30 °C water bath for 30 min, before harvesting and washing with 10 ml Sorbitol/Tris buffer (without β-mercaptoethanol). The spheroplasts were then resuspended in 1 ml of ice-cold NP buffer (1 M sorbitol, 50 mM NaCl, 10 mM Tris/HCl pH 7.5, 5 mM MgCl_2_, 1 mM CaCl_2_, 0.75% NP-40, 1 mM β-mercaptoethanol, 0.5 mM spermidine).

For MNase digestion, 1.5 U of MNase (Sigma-Aldrich) was added to each sample, and samples were incubated at 37 °C for 15 min. The reaction was stopped by the addition of 138 ul of Stop Buffer (5% SDS, 100 mM EDTA). Next, 40 µl of 10 mg/ml RNAse A (Thermo Fisher) was added and samples were incubated at 37 °C for 45 min, followed by the addition of 50 µl of 5 mg/ml Proteinase K (Thermo Fisher) and further incubation at 65 °C overnight.

The next morning, 360 ul of potassium acetate pH 5.5 was added to each sample, followed by incubation on ice for 10 min. DNA was then isolated from the supernatant using classical phenol-chloroform-isopropanol extraction and ethanol precipitation methods. Extracted DNA was then run on a 2% agarose gel, and bands corresponding to mononucleosomal DNA was recovered by gel extraction. DNA concentration was measured by Nanodrop spectrophotometer.

For library preparations, all samples, were processed using the NEBNext Ultra II library preparation kit for Illumina (NEB Cat. No. E7645L) using custom synthesized barcoded sequencing adapters provided by the Vienna BioCenter Core Facilities Next Generation Sequencing Facility. Following quality control by Agilent Fragment Analyzer and quantification by RT-qPCR using a Kapa Library Quantification Kit (Roche Cat. No. KK4903), libraries were sequenced at the Next Generation Sequencing Facility on an Illumina NovaSeq X. Alignments, quality control, and initial processing were performed using the nf-core/mnaseseq pipeline (https://github.com/nf-core/mnaseseq) with default parameters. Further analysis was performed using Deeptools (v3.3.1)^125^.

### ChIP-seq Library Generation and Analysis

*S. pombe* strains (100 ml) were grown to mid-exponential phase (OD_600_ ∼0.4-0.5) in YES, fixed with 1% formaldehyde for 10 min, quenched with 125 mM glycine for 10 min, then collected by centrifugation. Cells were washed once with ice-cold PBS, then resuspended in lysis buffer (50 mM HEPES-KOH, pH 7.5, 140 mM NaCl, 1 mM EDTA, 0.1% sodium deoxycholate, 1% Triton X-100, 1X Roche Complete Protease Inhibitors), and then lysed using acid-washed glass beads in a Precellys for 4×20 sec maximum speed, 1 min on ice between rounds. Lysate was then separated from the beads and then chromatin was sheared to a predominant final DNA length of ∼150-200 bp in a Covaris E220 ultrasonicator, and then clarified of unlysed cells and debris by centrifugation twice at 16,000 × *g* for 10 min. Prepared chromatin was then frozen at −70 °C pending use.

Chromatin aliquots were thawed on ice, then normalized based on starting culture density using additional lysis buffer for dilution. 10% of normalized and diluted chromatin were reserved as input controls. Either anti-Myc 9E10 (1µg; IMP Molecular Biology Service, AB_558470), anti-H3K36me3 (1µg; Abcam, ab9050), anti-H2BK120ub (1µg; Active Motif, Cat#39623) or anti-H2A.Z (2µg; Active Motif, Cat#39640) antibodies were then added to each culture and samples were incubated overnight at 4 °C with inversion. 25 µl of Protein A Dynabeads (Thermo Fisher Cat. No. 10001D) per sample were prepared by washing twice with PBS with 0.1% tween-20, once with lysis buffer, then resuspending in 100 µl of lysis buffer. Prepared beads were then combined with overnight chromatin samples and incubated for 4 hrs at 4 °C with inversion. Beads were then collected using the magnet and washed twice with lysis buffer, twice with high salt lysis buffer (50 mM HEPES-KOH, pH 7.5, 500 mM NaCl, 1 mM EDTA, 0.1% sodium deoxycholate, 1% Triton X-100), twice using lithium wash buffer (10 mM Tris-HCl, pH 8, 250 mM LiCl, 1 mM EDTA, 0.5% NP-40, 0.5% sodium deoxycholate), and once with TE. Beads were then resuspended in 100 µl of elution buffer (50 mM Tris-HCl, pH 8, 10 mM EDTA, 0.8% SDS) and transferred to PCR strips. Samples were then boiled for 10 min at 95 °C, then 65 °C overnight in a thermocycler. The next morning, proteinase K was added (final concentration 0.2 mg/mL) and incubated at 55 °C for 2 h. Samples were then cleaned up using a commercial kit (Zymo Research Cat. No. D5201). DNA concentration was measured by Nanodrop spectrophotometer.

For library preparations, all samples, including input, were processed using the NEBNext Ultra II library preparation kit for Illumina (NEB Cat. No. E7645L) using custom synthesized barcoded sequencing adapters provided by the Vienna BioCenter Core Facilities Next Generation Sequencing Facility. Following quality control by Agilent Fragment Analyzer and quantification by RT-qPCR using a Kapa Library Quantification Kit (Roche Cat. No. KK4903), libraries were sequenced at the Next Generation Sequencing Facility on an Illumina NovaSeq X. Alignments, quality control, and initial processing were performed using the nf-core/chipseq pipeline (https://github.com/nf-core/chipseq) with default parameters. Input-normalized bigWig files were subsequently generated using Deeptools (v3.3.1)^125^.

### AlphaFold predictions

We ran either AlphaFold-Multimer v2.2^81,126^ or AlphaFold3^127^ (https://alphafoldserver.com/) for all protein complex predictions. Of the five models generated, we used only the model ranked 1 for further processing and interpretation. We used the software UCSF ChimeraX (v1.8)^128^ for the visualization and superimposition of AlphaFold models. For the *in-silico* protein interaction screen, we used a previously established pipeline^129^ that integrated MMseqs and ColabFold^130^ (https://gitlab.com/BrenneckeLab/ht-colabfold).

### Multiple Sequence Alignments

Multiple sequence alignments were conducted for homologs of Hrp3 and Prf1 across 11 representative eukaryotic model species: *Schizosaccharomyces pombe, Saccharomyces cerevisiae, Neurospora crassa, Caenorhabditis elegans, Drosophila melanogaster, Danio rerio, Mus musculus, Homo sapiens, Physcomitrella patens, Arabidopsis thaliana,* and *Oryza sativa*. Alignments were performed using the Multiple Alignment using Fast Fourier Transform (MAFFT)^131^ algorithm. Homologs were identified as proteins clustering within the same orthogroup, based on Orthofinder^132^ (version 2.5.4) analysis performed with default settings.

### Quantification and Statistical Analysis

Unless otherwise noted, all statistical analyses were performed in R (v4.1.3) using the RStudio IDE (v2022.12.0+353) on x86_64-apple-darwin17.0 (64-bit) under macOS 13.6.3. All statistical tests used are stated in corresponding figure legends. Processing of NGS data, as well as all AlphaFold2 Multimer analyses, were performed using the CLIP cluster (https://clip.science).

