## supplemental figures and tables for "Nucleosome Positioning Shapes Cryptic Antisense Transcription"

**Supplementary Data**

**Figures S1-S9**

**Tables S1-S3**

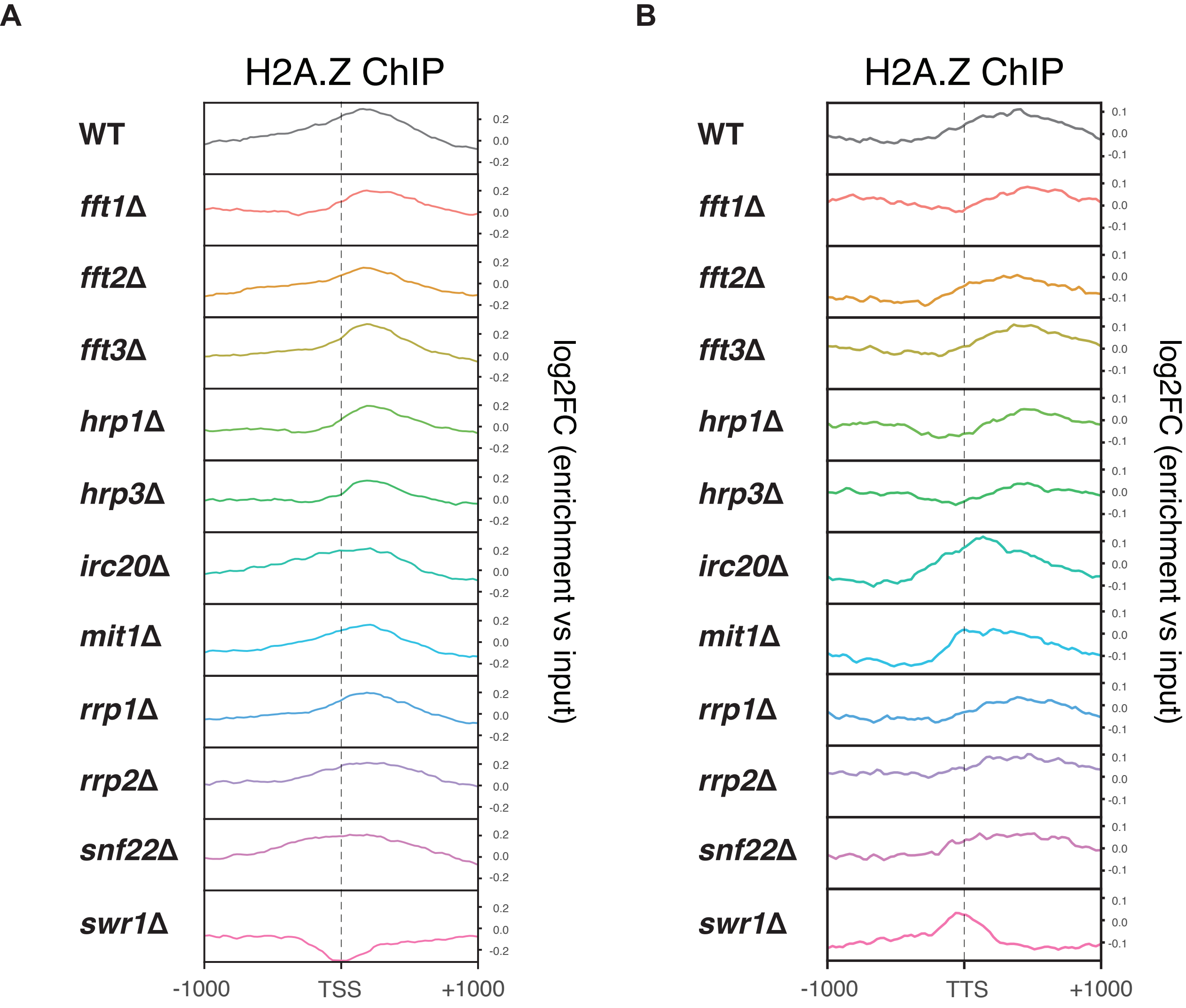
A
B
H2A.Z ChIP
H2A.Z ChIP
WT
fft1Δ
fft2Δ
fft3Δ
hrp1Δ
hrp3Δ
irc20Δ
mit1Δ
rrp1Δ
rrp2Δ
snf22Δ
swr1Δ
log2FC (enrichment vs input)
-1000
TSS
+1000
TTS
0.2
0.0
-0.2
0.1
-0.1

**Figure S1. H2A.Z Enrichment in 11 Nucleosome Remodeler Mutants in Fission Yeast**

(A) Metagene plots showing ChIP-seq coverage of H2A.Z relative to input across all protein-coding genes in WT, *fft1*Δ, *fft2*Δ, and *fft3*Δ, *hrp1*Δ and *hrp3*Δ, *irc20*Δ, *mit1*Δ, *rrp1*Δ, *rrp2*Δ, *snf22*Δ, and *swr1*Δ mutants. H2A.Z enrichment is plotted relative to the transcription start site (TSS), including 1 kb upstream and downstream.

(B) As in (A), but H2A.Z enrichment is plotted relative to the transcription termination site (TTS) instead.

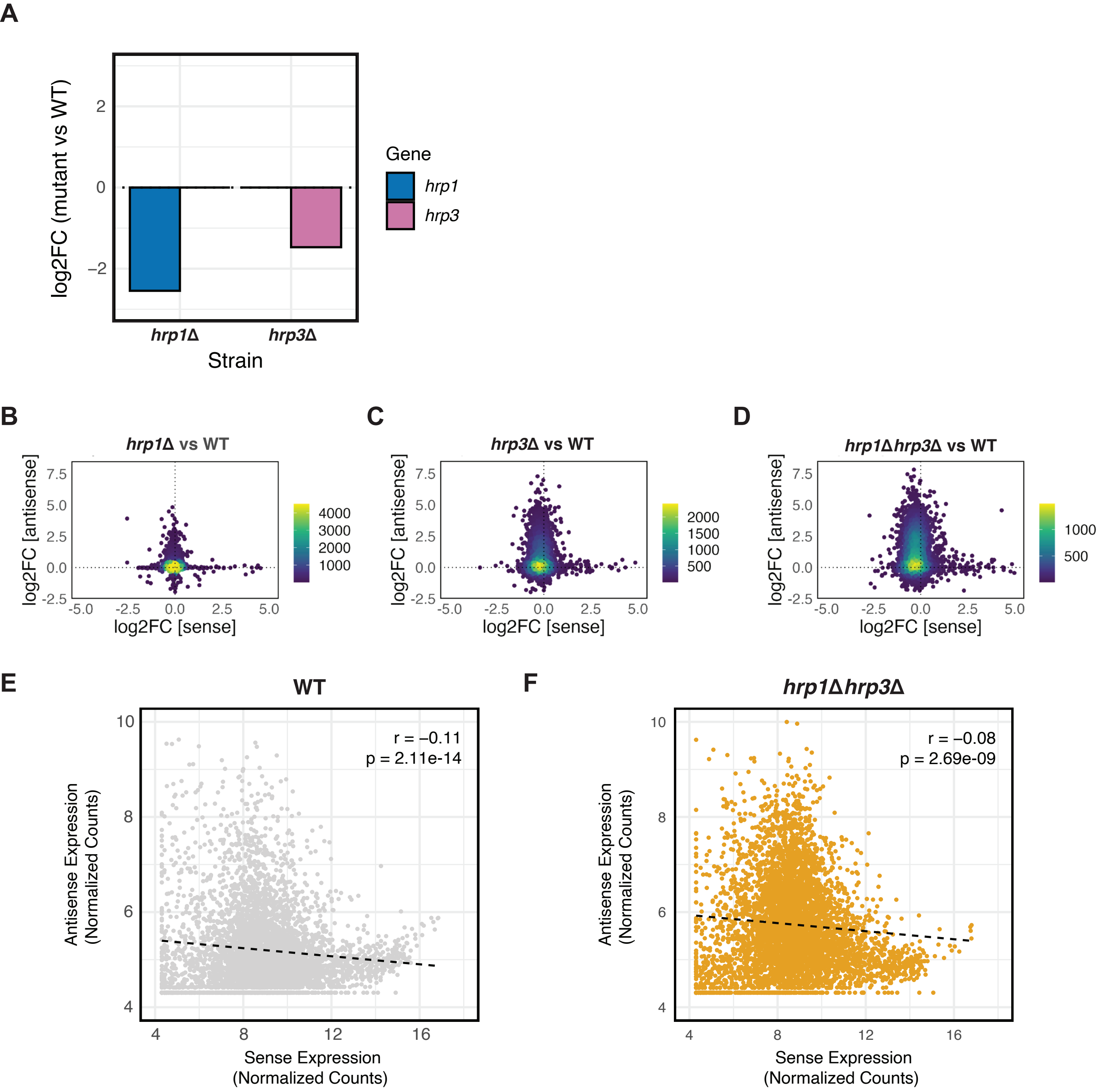
A
log2FC (mutant vs WT)
Gene
hrp1
hrp3
hrp1Δ
hrp3Δ
Strain
B
hrp1Δ vs WT
C
hrp3Δ vs WT
D
hrp1Δhrp3Δ vs WT
log2FC [antisense]
log2FC [sense]
E
WT
r = −0.11
p = 2.11e-14
F
hrp1Δhrp3Δ
r = −0.08
p = 2.69e-09
Antisense Expression
(Normalized Counts)
Sense Expression
(Normalized Counts)

**Figure S2. Analyses of Sense and Antisense Expression in *hrp1*Δ, *hrp3*Δ and *hrp1*Δ*hrp3*Δ.**

(A) Expression levels of *hrp1* and *hrp3* in *hrp3*Δ and *hrp1*Δ mutants, respectively. Expression is shown as log2 fold change (log2FC) relative to WT, based on mRNA-seq data. Values represent the average of three biological replicates.

(B) Density-colored scatterplots of mRNA-seq log2FC values for sense and antisense transcripts in *hrp1*Δ versus WT across all protein-coding genes. Data represents the average of three biological replicates.

(C) As in (B), but for *hrp3*Δ versus WT.

(D) As in (B), but for *hrp1*Δ*hrp3*Δ versus WT.

(E) Scatterplot comparing sense (x-axis) and antisense (y-axis) expression counts across all protein-coding genes in WT. Count data is derived from variance-stabilizing transformation (vst) of raw mRNA-seq counts. Pearson's correlation and linear regression analysis were performed. Data represents the average of three biological replicates.

(F) As in (E), but for the *hrp1*Δ*hrp3*Δ strain.

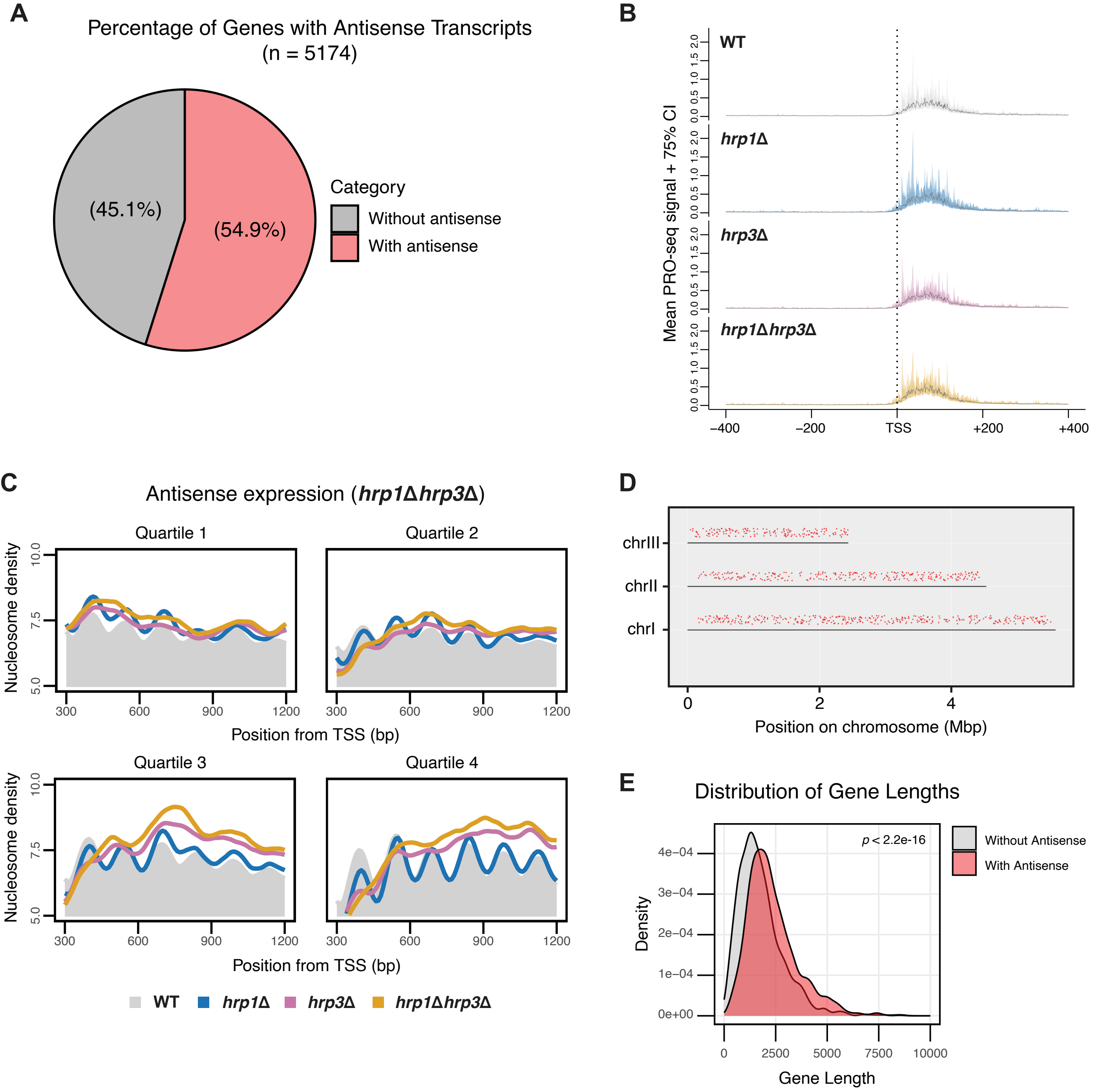

A
Percentage of Genes with Antisense Transcripts
(n = 5174)
(45.1%)
(54.9%)
Category
Without antisense
With antisense
B
Mean PRO-seq signal + 75% CI
WT
hrp1Δ
hrp3Δ
hrp1Δhrp3Δ
0.0 0.5 1.0 1.5 2.0
−400
−200
TSS
+200
+400
C
Antisense expression (hrp1Δhrp3Δ)
Quartile 1
Quartile 2
Quartile 3
Quartile 4
Nucleosome density
5.0
7.5
10.0
300
600
900
1200
Position from TSS (bp)
WT
hrp1Δ
hrp3Δ
hrp1Δhrp3Δ
D
chrIII
chrII
chrI
0
2
4
Position on chromosome (Mbp)
E
Distribution of Gene Lengths
p < 2.2e-16
Density
0e+00
1e−04
2e−04
3e−04
4e−04
0
2500
5000
7500
10000
Gene Length
Without Antisense
With Antisense

**Figure S3. Additional Analyses of Antisense Transcripts in *hrp1*Δ, *hrp3*Δ and *hrp1*Δ*hrp3*Δ.**

(A) Pie chart showing the proportion of all protein coding genes that have overlapping antisense transcripts (red) and those that do not (grey) in *S. pombe*. The percentages for each category are displayed on the chart.

(B) Metagene representation of mean PRO-seq signals (solid lines) with 75% confidence intervals (shaded regions) for WT, *hrp1Δ*, *hrp3Δ*, and *hrp1Δhrp3Δ* mutants centered on the TSS of the all protein-coding genes in fission yeast. Profiles include 400 bp upstream and downstream of the TSS. Data represent the mean signal from two biological replicates.

(C) Metaplots of normalized nucleosome density from MNase-seq within the gene body (300 bp to 1200 bp from the TSS) of all protein-coding genes for WT, *hrp1Δ*, *hrp3Δ*, and *hrp1Δhrp3Δ*. Data are stratified into four quartiles based on antisense expression levels (lowest to highest) in the *hrp1Δhrp3Δ* mutant. The WT profile is shown in solid grey, and the plotted data represent the average signal from two biological replicates.

(D) Distribution of antisense expression across chromosomes I, II, and III in the *hrp1Δhrp3Δ* mutant. Each red dot represents a gene with an antisense transcript.

(E) The density plot shows the distribution of gene lengths for two groups: genes with antisense transcripts (red) and genes without antisense transcripts (grey). The Wilcoxon rank sum test was used to compare the two distributions for statistical significance.

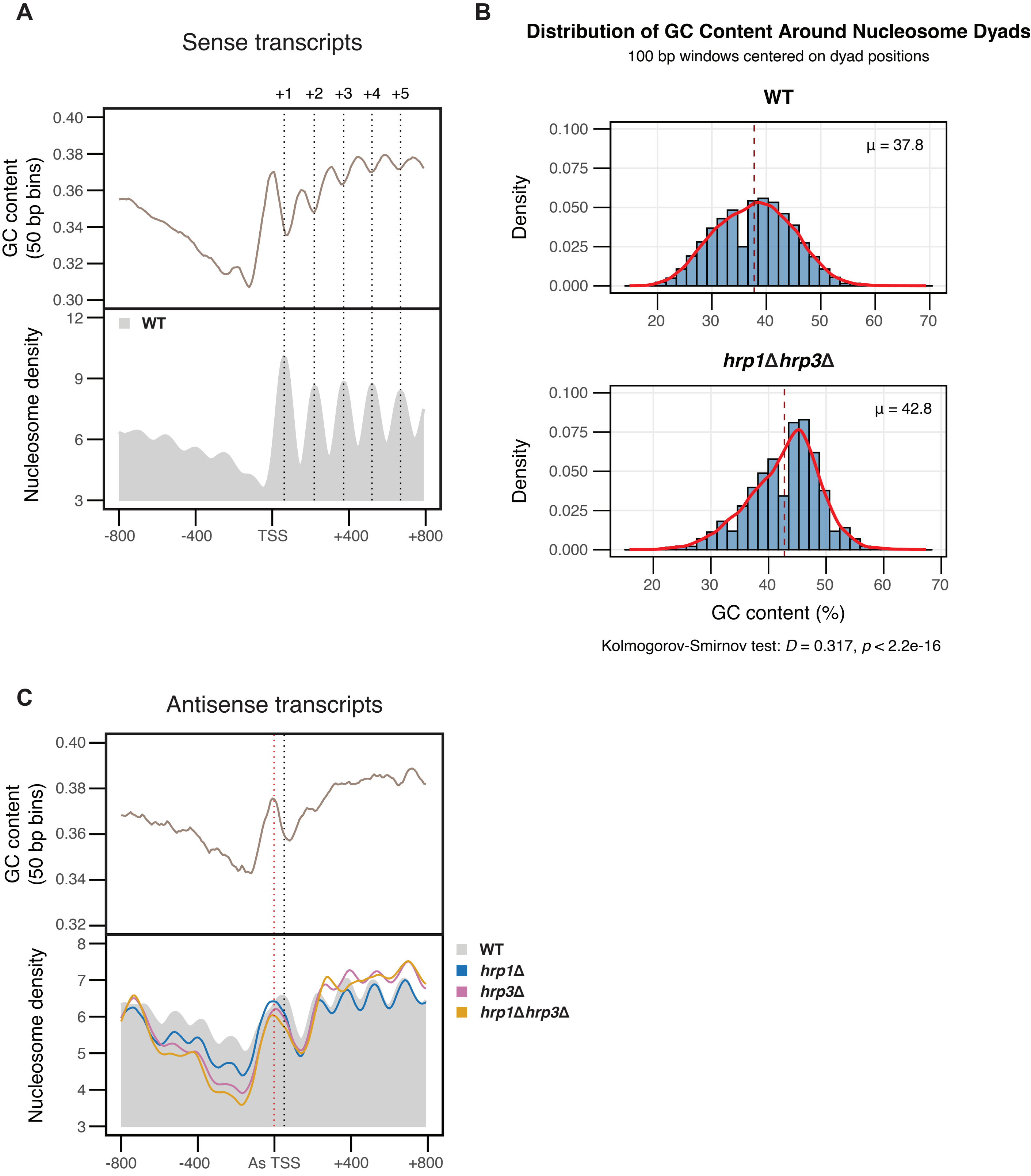
A
Sense transcripts
+1
+2
+3
+4
+5
GC content
(50 bp bins)
0.40
0.38
0.36
0.34
0.32
0.30
Nucleosome density
12
9
6
3
WT
-800
-400
TSS
+400
+800
B
Distribution of GC Content Around Nucleosome Dyads
100 bp windows centered on dyad positions
WT
Density
μ = 37.8
0.100
0.075
0.050
0.025
0.000
20
30
40
50
60
70
hrp1Δhrp3Δ
μ = 42.8
GC content (%)
Kolmogorov-Smirnov test: D = 0.317, p < 2.2e-16
C
Antisense transcripts
GC content
(50 bp bins)
0.40
0.38
0.36
0.34
0.32
Nucleosome density
8
7
6
5
4
3
WT
hrp1Δ
hrp3Δ
hrp1Δhrp3Δ
-800
-400
As TSS
+400
+800

**Figure S4. Loss of *hrp3* is Associated with Depletion of Nucleosome Occupancy over AT-rich Regions**

(A) (Top) Metaplot of genomic GC content (50 bp bins) across the TSS of all protein coding genes (Bottom) Metaplot of normalized nucleosome density from MNase-seq across the same region. Dotted lines indicate positions of nucleosomes downstream of the TSS. Profiles include 800 bp upstream and downstream of the TSS.

(B) Density histograms of GC content within 100 bp windows centered on all nucleosome dyads in WT and *hrp1Δhrp3Δ*. The dotted line indicates the mean GC content (μ). The Kolmogorov-Smirnov test was used to compare the two distributions for statistical significance.

(C) (Top) Metaplot of genomic GC content (50 bp bins) across the antisense (As) TSS of 4,678 antisense transcripts. (Bottom) Metaplot of normalized nucleosome density from MNase-seq across the same region (as shown in Fig. 3C). The black dotted line indicates the position of the well-positioned nucleosome in WT, while the red dotted line indicates the shifted position of this nucleosome in the mutants. Profiles include 800 bp upstream and downstream of the As-TSS.

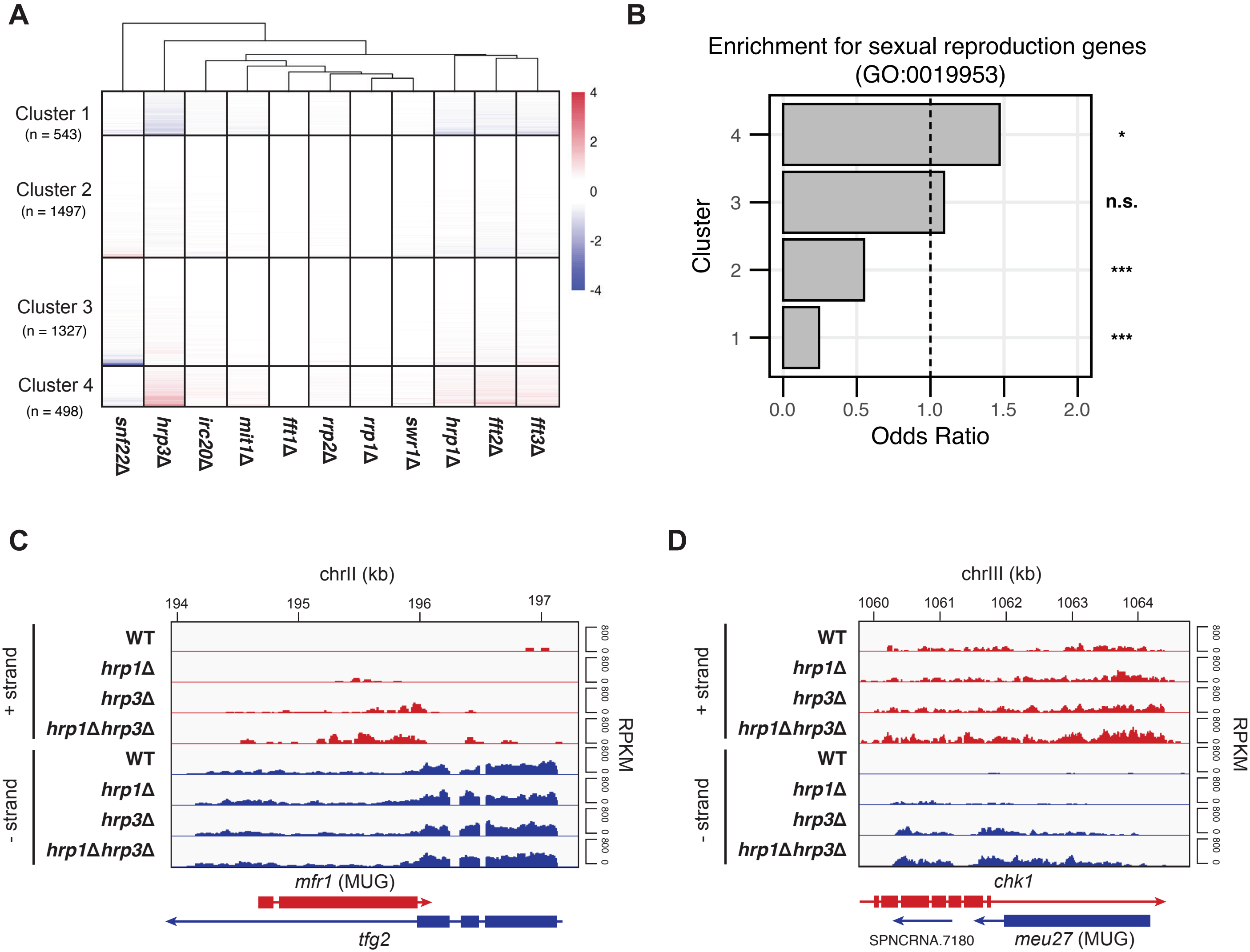

A
Cluster 1
(n = 543)
Cluster 2
(n = 1497)
Cluster 3
(n = 1327)
Cluster 4
(n = 498)
4
2
0
-2
-4
snf22Δ
hrp3Δ
irc20Δ
mit1Δ
fft1Δ
rrp2Δ
rrp1Δ
swr1Δ
hrp1Δ
fft2Δ
fft3Δ
B
Enrichment for sexual reproduction genes (GO:0019953)
Cluster
4
3
2
1
*
n.s.
***
***
0.0
0.5
1.0
1.5
2.0
Odds Ratio
C
chrII (kb)
194
195
196
197
+ strand
- strand
WT
hrp1Δ
hrp3Δ
hrp1Δhrp3Δ
RPKM
800
0
mfr1 (MUG)
tfg2
D
chrIII (kb)
1060
1061
1062
1063
1064
chk1
SPNCRNA.7180
meu27 (MUG)

**Figure S5. Further Analyses on MUGs and Convergent Nested Genes in Fission Yeast.**

(A) Heatmap of significantly differentially expressed protein-coding genes (adj p-value < 0.05, DESeq2) in 11 nucleosome remodeler deletion mutants relative to WT, based on mRNA-seq data. The color scale represents log2 fold change (log2FC) in gene expression for each mutant compared to WT. Genes are stratified into four clusters based on their expression patterns.

(B) Enrichment analysis of the differentially expressed genes from (A) across the four clusters for the sexual reproduction GO term (GO:0019953). The dashed line indicates an odds ratio of 1. Statistical analysis was performed using Fisher's exact test. Asterisks indicate statistical significance: $p < 0.05$ (*), $p < 0.01$ (**), $p < 0.001$ (***).

(C) Genome browser track of mRNA-seq data for the *mfr1*/*tfg2* nested gene pairs in WT, *hrp1*Δ, *hrp3*Δ *and hrp1*Δ*hrp3*Δ. Tracks are separated into the + strand (red) and – strand (blue), with values shown in RPKM. The MUG within the pair is denoted in brackets.

(D) As in (C), but for the *chk1*/ *meu27* nested gene pair.

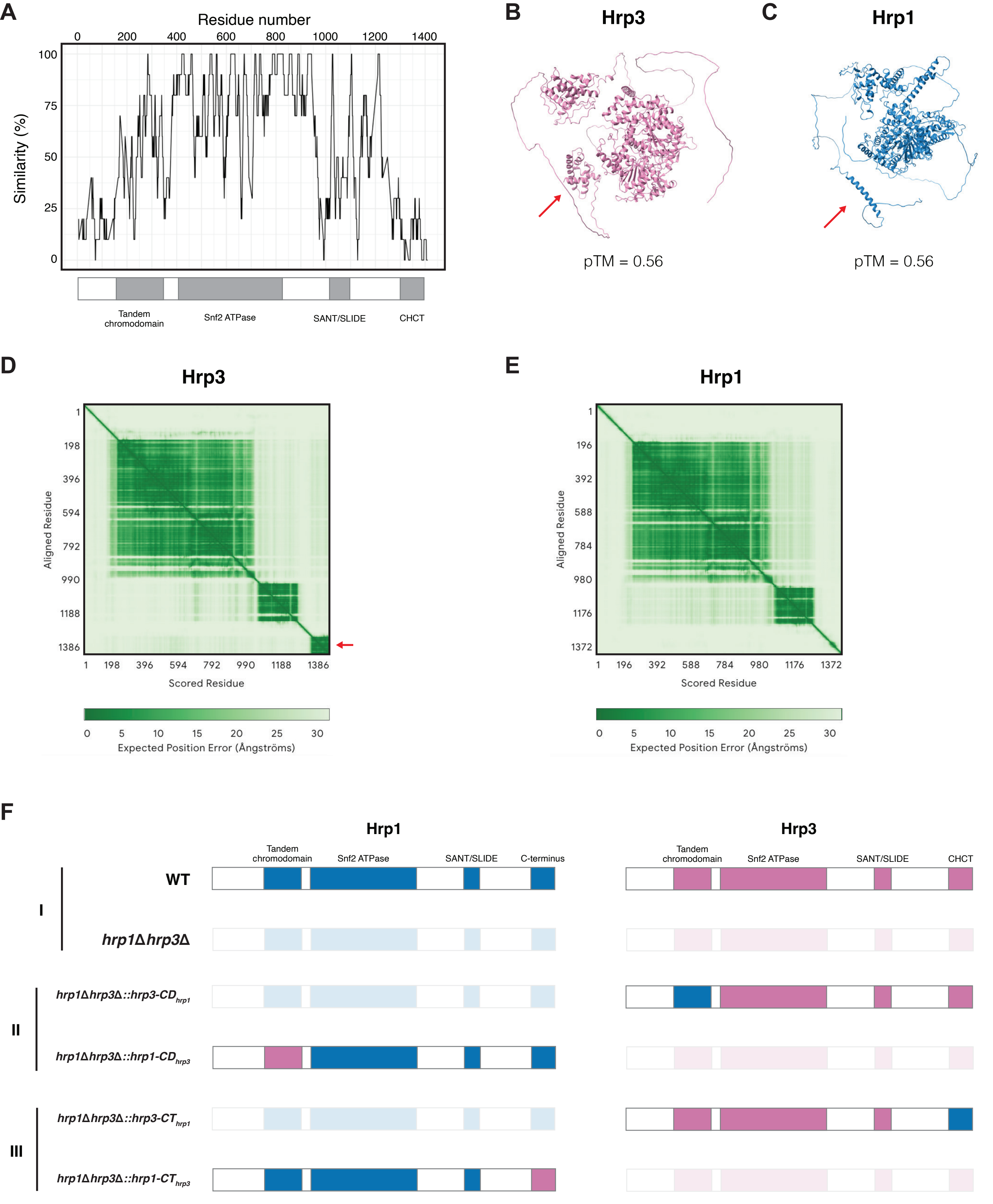
A
Residue number
0 200 400 600 800 1000 1200 1400
Similarity (%)
100 75 50 25 0
Tandem chromodomain
Snf2 ATPase
SANT/SLIDE
CHCT
B
Hrp3
pTM = 0.56
C
Hrp1
pTM = 0.56
D
Hrp3
Aligned Residue
1 198 396 594 792 990 1188 1386
Scored Residue
Expected Position Error (Ångströms)
0 5 10 15 20 25 30
E
Hrp1
Aligned Residue
1 196 392 588 784 980 1176 1372
Scored Residue
Expected Position Error (Ångströms)
0 5 10 15 20 25 30
F
Hrp1
Hrp3
Tandem chromodomain
Snf2 ATPase
SANT/SLIDE
C-terminus
CHCT
I
WT
hrp1Δhrp3Δ
II
hrp1Δhrp3Δ::hrp3-CD hrp1
hrp1Δhrp3Δ::hrp1-CD hrp3
III
hrp1Δhrp3Δ::hrp3-CT hrp1
hrp1Δhrp3Δ::hrp1-CT hrp3

**Figure S6. Comparative Domain Analysis of Hrp1 and Hrp3.**

(A) Sliding window analysis of protein sequence similarity between Hrp1 and Hrp3. Sequence similarity was calculated using a sliding window of 25 residues after multiple sequence alignment (MSA) with the msa R package (default settings). Each point represents the similarity score for a specific window, plotted against the central residue position. Gaps in the alignment were excluded to ensure accurate similarity calculations. Protein domains are annotated below the plot.

(B) AlphaFold3-predicted structure of Hrp3. The red arrow marks the C-terminus. The pTM value represents the Predicted Template Modelling (pTM) score.

(C) AlphaFold3-predicted structure of Hrp1. The red arrow indicates the CHCT domain. The pTM value represents the Predicted Template Modelling (pTM) score.

(D) Predicted Aligned Error (PAE) plot from AlphaFold3 for Hrp3, showing the expected position error for each residue.

(E) PAE plot for Hrp1, as in (D). The red arrow highlights the expected positional error for the CHCT domain.

(F) Schematic representation of protein domain configurations in Hrp1 and Hrp3 for wild-type (WT) and mutant strains. Mutants include *hrp1Δhrp3Δ*, *hrp1Δhrp3Δ::hrp3-CD*$_{hrp1}$, *hrp1Δhrp3Δ::hrp1-CD*$_{hrp3}$, *hrp1Δhrp3Δ::hrp3-CT*$_{hrp1}$, and *hrp1Δhrp3Δ::hrp1-CT*$_{hrp3}$. CD stands for chromodomain, while CT stands for C-terminus. Domain swaps are indicated by color changes, and greyed-out proteins represent their absence in the respective mutants.

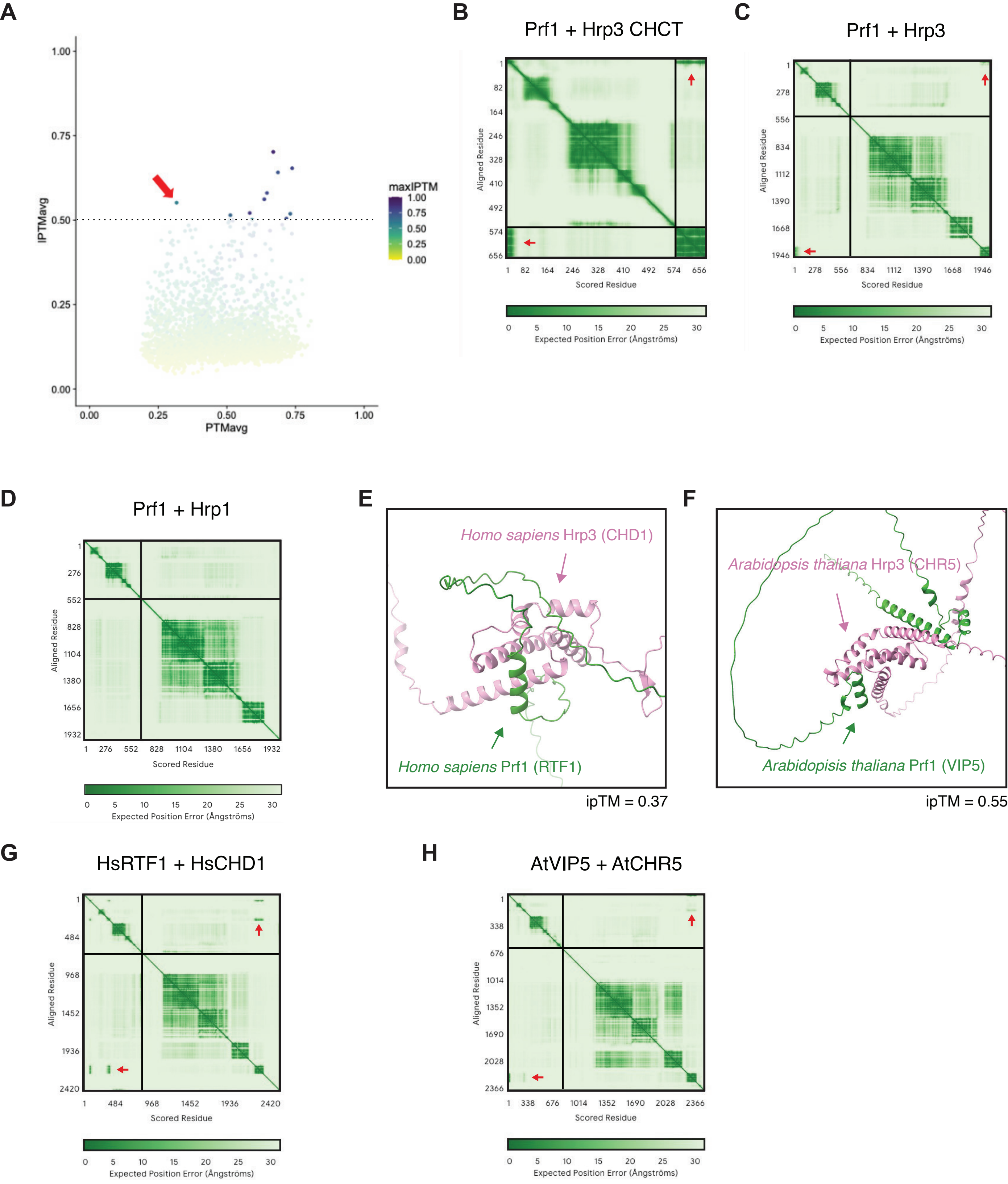

A
IPTMavg
PTMavg
1.00
0.75
0.50
0.25
0.00
maxIPTM
B
Prf1 + Hrp3 CHCT
C
Prf1 + Hrp3
D
Prf1 + Hrp1
Aligned Residue
Scored Residue
Expected Position Error (Ångströms)
0
5
10
15
20
25
30
E
Homo sapiens Hrp3 (CHD1)
Homo sapiens Prf1 (RTF1)
ipTM = 0.37
F
Arabidopsis thaliana Hrp3 (CHR5)
Arabidopisis thaliana Prf1 (VIP5)
ipTM = 0.55
G
HsRTF1 + HsCHD1
H
AtVIP5 + AtCHR5

**Figure S7. AlphaFold-predicted Interactions Between Prf1 and the Hrp3 CHCT Domain Across Species.**

(A) Scatterplot of the average Interface Predicted Template Modelling (ipTM) score versus the average Predicted Template Modelling (pTM) score from an *in-silico* screen using AlphaFold2 Multimer. The screen was performed on the Hrp3 CHCT domain with a set of 2,692 genes known to localize to the *S. pombe* nucleus. The dotted line indicates an average ipTM score cutoff of 0.5. The red arrow highlights Prf1.

(B) Predicted Aligned Error (PAE) plot from AlphaFold3 showing the predicted binding between Prf1 and the CHCT domain of Hrp3. Red arrows indicate the predicted interface with low expected positional error between the N-terminus of Prf1 and the CHCT domain.

(C) As in (B), but showing the predicted binding between Prf1 and full-length Hrp3. Red arrows indicate the predicted interface with low expected positional error between the N-terminus of Prf1 and the C-terminus of Hrp3.

(D) As in (B), but showing the predicted binding between Prf1 and full-length Hrp1.

(E) AlphaFold3-predicted structure of the interaction between the N-terminus of *Homo sapiens* Prf1 (RTF1) and the CHCT domain of *Homo sapiens* Hrp3 (CHD1).

(F) As in (E), but showing the interaction between the N-terminus of *Arabidopsis thaliana* Prf1 (VIP5) and the CHCT domain of *Arabidopsis thaliana* Hrp3 (CHR5).

(G) Predicted Aligned Error (PAE) plot from AlphaFold3 showing the predicted binding between *Homo sapiens* RTF1 and CHD1. Red arrows indicate the predicted interface with low expected positional error between the N-terminus of RTF1 and the C-terminus of CHD1.

(H) As in (F), but showing the predicted binding between *Arabidopsis thaliana* VIP5 and CHR5. Red arrows indicate the predicted interface with low expected positional error between the N-terminus of VIP5 and the C-terminus of CHR5.

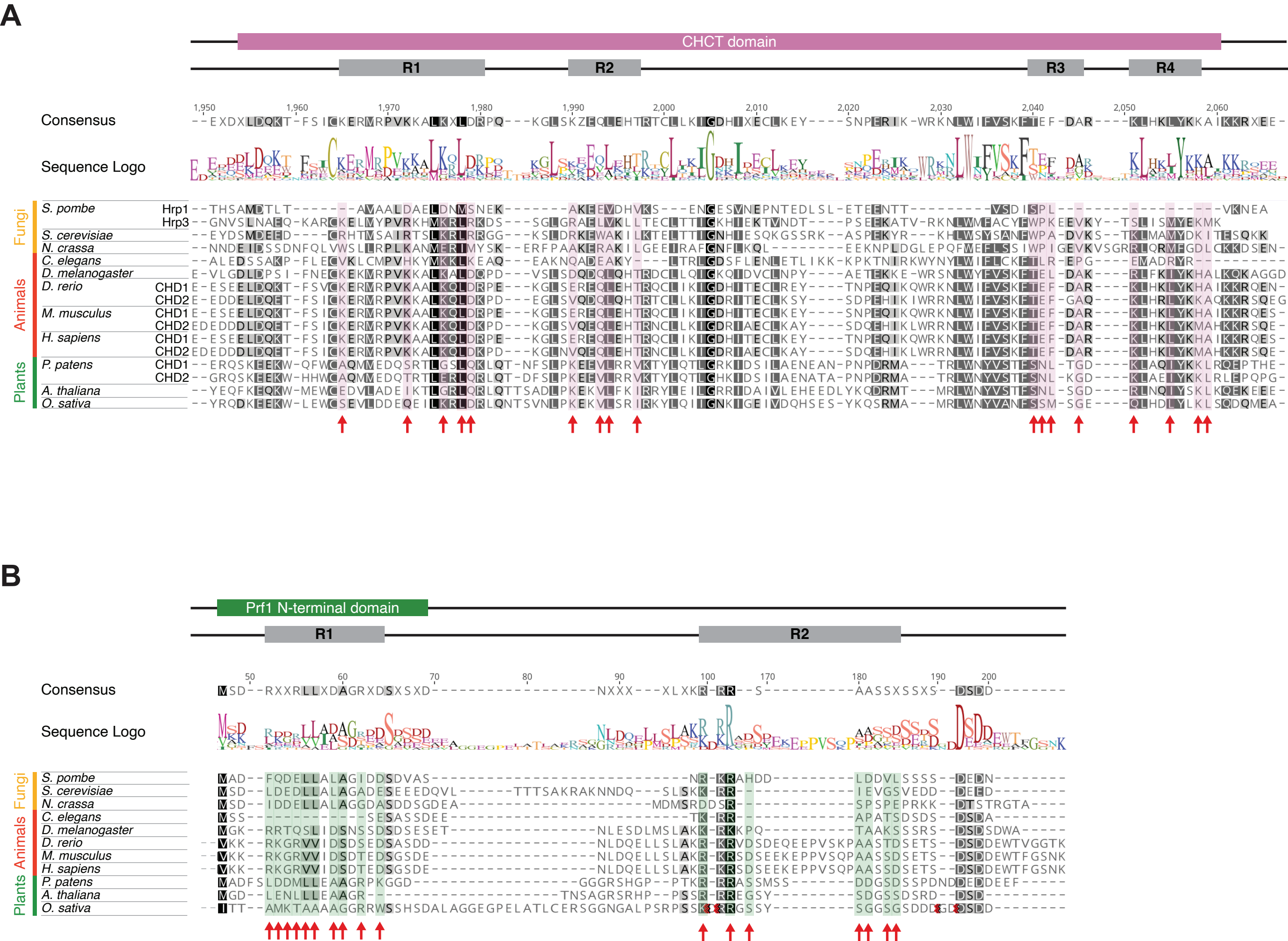

A
CHCT domain
R1
R2
R3
R4
Consensus
Sequence Logo
Fungi
Animals
Plants
S. pombe
Hrp1
Hrp3
S. cerevisiae
N. crassa
C. elegans
D. melanogaster
D. rerio
CHD1
CHD2
M. musculus
CHD1
CHD2
H. sapiens
CHD1
CHD2
P. patens
CHD1
CHD2
A. thaliana
O. sativa
B
Prf1 N-terminal domain
R1
R2
Consensus
Sequence Logo
Fungi
Animals
Plants
S. pombe
S. cerevisiae
N. crassa
C. elegans
D. melanogaster
D. rerio
M. musculus
H. sapiens
P. patens
A. thaliana
O. sativa

**Figure S8. Interacting Residues Between Hrp3 and Prf1 are Conserved Across Eukaryotes.**

(A) Multiple sequence alignment (MSA) of the CHCT domain of Hrp3 across 11 eukaryotic species, performed using Multiple Alignment using Fast Fourier Transform (MAFFT). For species with two paralogs, both copies are included. Residues are shaded in increasing intensities of black based on their degree of conservation across species. Residues predicted by AlphaFold3 to interact with Prf1 are annotated with red arrows and highlighted in pink, corresponding to four regions (R1, R2, R3, and R4). Hrp1 is included as a comparison, despite lacking the CHCT domain. Species used in the analysis were *Schizosaccharomyces pombe, Saccharomyces cerevisiae, Neurospora crassa, Caenorhabditis elegans, Drosophila melanogaster, Danio rerio, Mus musculus, Homo sapiens, Physcomitrella patens, Arabidopsis thaliana,* and *Oryza sativa*.

(B) As in (A), but showing the MSA results for the N-terminal region of Prf1. Residues predicted by AlphaFold3 to interact with Hrp3 are annotated with red arrows and highlighted in green, corresponding to two regions (R1 and R2).

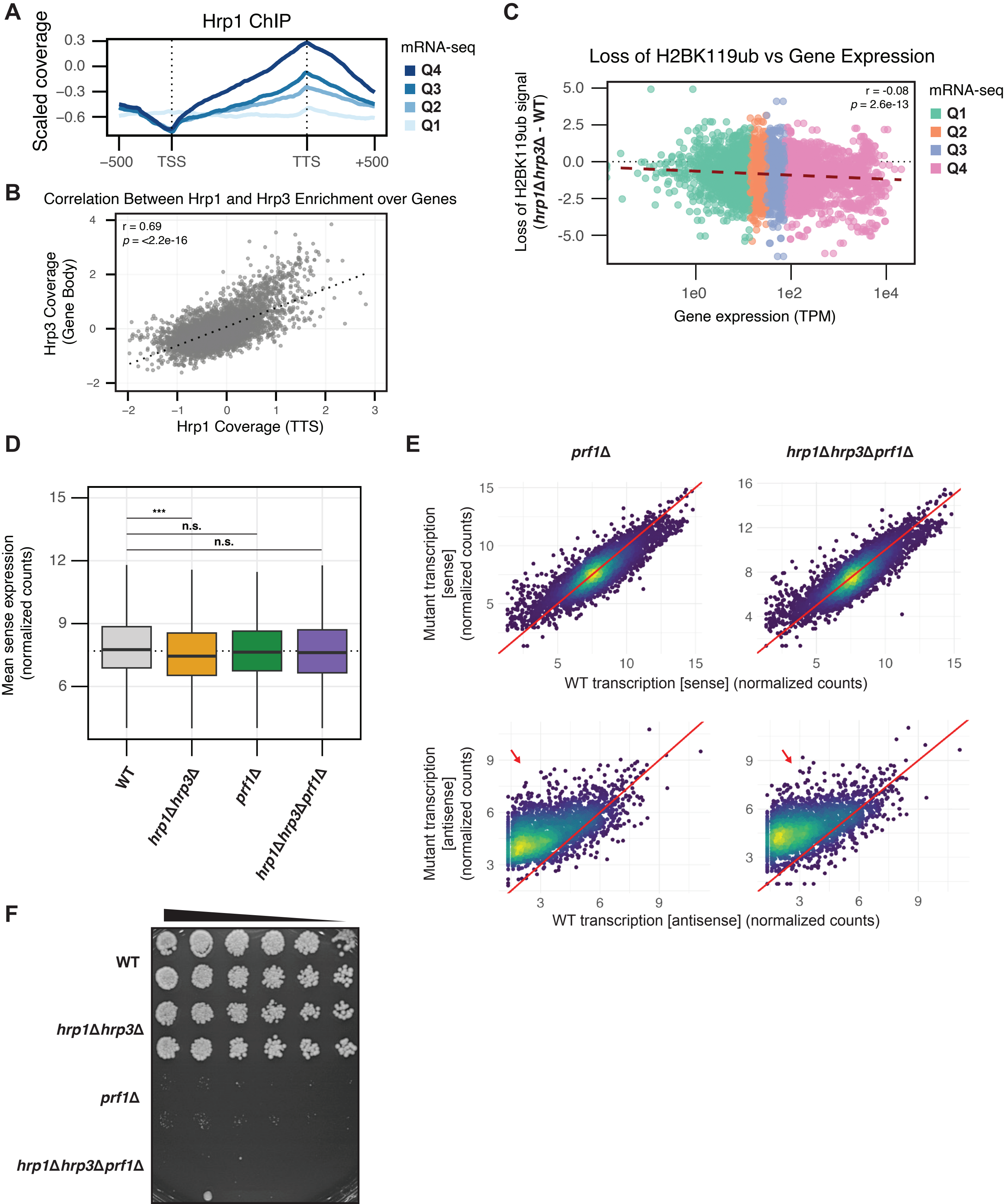
A
Hrp1 ChIP
Scaled coverage
0.3
0.0
−0.3
−0.6
−500
TSS
TTS
+500
mRNA-seq
Q4
Q3
Q2
Q1
B
Correlation Between Hrp1 and Hrp3 Enrichment over Genes
r = 0.69
p = <2.2e-16
Hrp3 Coverage (Gene Body)
Hrp1 Coverage (TTS)
C
Loss of H2BK119ub vs Gene Expression
r = -0.08
p = 2.6e-13
Loss of H2BK119ub signal (hrp1Δ hrp3Δ - WT)
Gene expression (TPM)
1e0
1e2
1e4
mRNA-seq
Q1
Q2
Q3
Q4
D
Mean sense expression (normalized counts)
***
n.s.
n.s.
WT
hrp1Δhrp3Δ
prf1Δ
hrp1Δhrp3Δprf1Δ
E
prf1Δ
hrp1Δhrp3Δprf1Δ
Mutant transcription [sense] (normalized counts)
WT transcription [sense] (normalized counts)
Mutant transcription [antisense] (normalized counts)
WT transcription [antisense] (normalized counts)
F
WT
hrp1Δhrp3Δ
prf1Δ
hrp1Δhrp3Δprf1Δ

**Figure S9. Additional Analyses of *prf1* mutants.**

(A) Metagene plot of the ChIP-seq coverage relative to input for Hrp1-myc in WT across all protein coding genes. Genes were divided into four quartiles based on expression in WT. Quartiles are numbered by expression level, with Q1 being the lowest and Q4 being the highest.

(B) Scatterplot showing the correlation between the ChIP-seq coverage relative to input for Hrp3 in the gene body versus the ChIP-seq coverage relative to input for Hrp1 at the TTS for all protein coding genes. Each point represents a single gene, with the x-axis corresponding to the mean Hrp1 signal in a 500 bp region centered on the TTS and the y-axis corresponding to the mean Hrp3 signal across the gene body (from TSS to TTS). The black dotted line represents the linear regression fit. The Pearson correlation coefficient (r) and p-value (p) are displayed on the plot.

(C) Scatterplot showing the correlation between the loss of H2BK119ub in *hrp1*Δ*hrp3*Δ*prf1*Δ versus WT and gene expression in WT across all protein-coding genes. The dashed line represents the linear regression. Points are color-coded by quartiles of gene expression. The dotted line at $y = 0$ indicates no loss of H2Bub signal. The Spearman correlation coefficient and p-value are displayed on the plot.

(D) Boxplot of variance-stabilizing transformation (vst)-normalized sense expression counts for WT, *hrp1*Δ*hrp3*Δ, *prf1*Δ, and *hrp1*Δ*hrp3*Δ*prf1*Δ. The dotted line represents the median sense expression in WT. Statistical analysis was performed using ANOVA with Tukey's Honestly Significant Difference (HSD) test. Asterisks indicate statistical significance: $p < 0.05$ (*), $p < 0.01$ (**), $p < 0.001$ (***).

(E) Density-colored scatterplots comparing transcription in mutants (*prf1*Δ and *hrp1*Δ*hrp3*Δ*prf1*Δ) to WT for sense transcripts (top row) and antisense transcripts (bottom row) across all protein-coding genes. Count data are derived from vst-transformed raw mRNA-seq counts. Red arrows highlight elevated antisense transcription in mutants relative to WT.

(F) Spot growth assay of WT, *hrp1*Δ*hrp3*Δ*, prf1*Δ and *hrp1*Δ*hrp3*Δ*prf1*Δ on YES agar. Spots represent serial 1:2 dilutions of each strain.

### Supplementary Tables

**Table S1**. Early, middle and late MUGs upregulated in *hrp3*Δ.

| Gene ID | Name | Description |
|---|---|---|
| | | **Early MUGs** |
| SPAC17A5.18c | rec25 | meiotic recombination protein Rec25 |
| SPAC25G10.04c | rec10 | meiotic recombination protein Rec10 |
| SPAC6C3.05 | meu43 | Schizosaccharomyces specific protein Meu43 |
| SPAC8E11.03c | dmc1 | RecA family ATPase Dmc1 |
| SPBC21B10.12 | rec6 | meiotic recombination protein Rec6 |
| SPBC31F10.05 | mug37 | conserved fungal protein, OB fold, predicted role in DNA metabolism |
| SPCC1620.04c | fzr3 | meiotic fizzy-related APC coactivator Fzr3 |
| | | **Middle MUGs** |
| SPAC1250.02 | mug95 | Schizosaccharomyces specific protein Mug95 |
| SPAC1610.03c | crp79 | poly(A) binding protein Crp79 |
| SPAC16A10.08c | mug74 | Schizosaccharomyces specific protein Mug74 |
| SPAC16E8.05c | | Schizosaccharomyces specific protein Mde1 |
| SPAC1A6.06c | meu31 | Schizosaccharomyces specific protein Meu31 |
| SPAC212.02 | | Schizosaccharomyces pombe specific protein |
| SPAC22F3.02 | atf31 | DNA-binding transcription factor Atf31 |
| SPAC22F3.04 | mug62 | implicated in acetyl-CoA biosynthesis and diacylglycerol metabolism |
| SPAC24C9.15c | spn5 | meiotic septin Spn5 |
| SPAC25H1.09 | mde5 | alpha-amylase homolog Mde5 |
| SPAC343.07 | mug28 | RNA-binding protein Mug28, implicated in mRNA processing |
| SPAC4G9.05 | mpf1 | cytoplasmic meiotic pumilio family RNA-binding protein Mpf1 |
| SPAC6B12.06c | rrg9 | mitochondrial ribosome assembly protein Rrg9 |
| SPAC6C3.07 | mug68 | Schizosaccharomyces specific protein Mug68 |
| SPAC6G10.06 | tda3 | FAD-dependent amino acid oxidase involved in late endosome to Golgi transport Tda3 |
| SPAC8F11.05c | mug130 | Schizosaccharomyces specific protein Mug130 |
| SPAC977.06 | | S. pombe specific DUF999 family protein 3 |

| | | |
|---|---|---|
| SPAPB17E12.09 | | conserved protein, expressed during meiotic cell cycle, possibly related to metazoan RAB11FIP4 family |
| SPBC1198.12 | mfr1 | meiotic APC activator Mfr1 |
| SPBC146.11c | mug97 | meiotically upregulated gene Mug97 |
| SPBC1685.06 | cid11 | poly(A) polymerase Cid11, terminal uridylyl transferase |
| SPBC16A3.13 | meu7 | alpha-amylase homolog Aah4 |
| SPBC1778.04 | spo6 | Spo4-Spo6 kinase complex regulatory subunit Spo6 |
| SPBC1861.06c | mug131 | UPF0300 family protein 4 |
| SPBC21D10.08c | | Schizosaccharomyces specific protein |
| SPBC27.03 | meu25 | Schizosaccharomyces specific protein Meu25 |
| SPBC32H8.06 | mug93 | Paqosome core subunit, TPR repeat protein, human RPAP3 ortholog |
| SPBC428.07 | meu6 | pleckstrin homology domain protein Meu6 |
| SPBC4C3.08 | otg2 | alpha-1,3-galactosyltransferase Otg2 |
| SPBC56F2.03 | arp10 | dynactin complex actin-like protein Arp10 |
| SPBC8D2.19 | mde3 | serine/threonine protein kinase, meiotic, STKc MAK-like Mde3 |
| SPCC11E10.09c | | alpha-amylase homolog |
| SPCC1235.13 | ght6 | plasma membrane glucose/fructose:proton symporter Ght6 |
| SPCC1259.14c | meu27 | UPF0300 family protein 5 |
| SPCC1682.12c | ubp16 | ubiquitin C-terminal hydrolase Ubp16 |
| SPCC188.12 | spn6 | meiotic (sporulation) septin Spn6 |
| SPCC1919.11 | mug137 | BAR adaptor protein, human endophilin A3-like |
| SPCC31H12.06 | mug111 | major facilitator family transmembrane transporter Mug111 |
| SPCC320.07c | mde7 | RNA-binding protein Mde7 |
| SPCC417.12 | | carboxylesterase, type B family protein |
| **Late MUGs** | | |
| SPAC15E1.02c | | DUF1761 transmembrane protein family, implicated in phosphate metabolism, stress response, sporulation |
| SPAC186.02c | | hydroxyacid dehydrogenase, implicated in cellular detoxification |
| SPAC22G7.11c | cum1 | Con-6 family conserved fungal protein |
| SPAC23D3.05c | | alcohol dehydrogenase pseudogene |
| SPAC2F7.06c | pol4 | DNA polymerase X family |
| SPAC3C7.02c | pil2 | meiotic eisosome BAR domain protein Pil2 |

| | | |
|---|---|---|
| SPAC4F10.17 | | plasma membrane and mitochondrial outer membrane, implicated in cell integrity signaling, conserved fungal protein |
| SPAC750.02c | | transmembrane transporter |
| SPAC869.06c | hry1 | HHE domain cation binding protein, implicated in the repair of iron-sulfur clusters damaged by oxidative and nitrosative stress |
| SPAC869.07c | mel1 | alpha-galactosidase, melibiase |
| SPAC869.08 | pcm2 | protein-L-isoaspartate O-methyltransferase Pcm2 |
| SPAC869.09 | | Con-6 family conserved fungal protein |
| SPAPB1A11.03 | | FMN-dependent alpha-hydroxy acid dehydrogenase, probable lactate dehydrogenase |
| SPAPB8E5.10 | min8 | mitochondrial respiratory complex/ATP synthase complex assembly protein Mra1, meiosis specific splicing in fission yeast |
| SPBC685.03 | | methyltransferase, sterol related |
| SPBC725.06c | ppk31 | serine/threonine protein kinase Ppk31 |
| SPBP4G3.03 | fub2 | PI31 proteasome regulator Fub2 |
| SPCC663.14c | trp663 | plasma membrane TRP-like ion channel |
| SPCC757.02c | | dehydrogenase |

**Table S2**. Candidates for predicted interaction with the CHCT domain of Hrp3 with ipTM scores greater than 0.5.

| Rank | Gene ID | NAME | IPTMavg | PTMavg | Description |
|---|---|---|---|---|---|
| 1 | SPCC830.11c | Fap7 | 0.70198 | 0.6692 | Ribosome assembly chaperone for Rps14 |
| 2 | SPAC26A3.17c | Rmt2 | 0.6536 | 0.7382 | N-methyltransferase |
| 3 | SPBC713.05 | Wdr83 | 0.6412 | 0.6864 | Gpl1-Gih35-Wdr83 complex WD repeat subunit Wdr83 |
| 4 | SPAC22F3.05c | Alp41 | 0.5804 | 0.6466 | GTP-binding protein involved in beta-tubulin folding Alp41 |
| 5 | SPBC18E5.08 | | 0.5618 | 0.6366 | N-acetyltransferase |
| 6 | SPBC651.09c | Prf1 | 0.5516 | 0.317 | RNA polymerase II associated Paf1 complex |
| 7 | SPBC685.04c | Aps2 | 0.521 | 0.5838 | AP-2 adaptor complex sigma subunit Aps2 |
| 8 | SPCC63.06 | | 0.5188 | 0.7314 | WD repeat protein, human WDR89 family |
| 9 | SPAC1B3.12c | Rpb10 | 0.5146 | 0.513 | DNA-directed RNA polymerase I, II, and III subunit Rpb10 |
| 10 | SPCC1672.01 | | 0.5046 | 0.7174 | histidinol-phosphatase |
| 11 | SPAC1782.08c | Rex3 | 0.5008 | 0.5926 | exonuclease Rex3 |
| 12 | SPCC1393.14 | Ten1 | 0.4982 | 0.5216 | telomere cap complex subunit Ten1 |

**Table S3.** *Schizosaccharomyces pombe* strains used in this study.

| Strain name | Genotype |
|---|---|
| YZH_001 | h+ ade6-M216 ura4-D18 leu1-32 |
| YZH_002 | h- ade6-M216 ura4-D18 leu1-32 |
| YZH_069 | h- ade6-M216 ura4-D18 leu1-32 hrp1-13xmyc-kanMX6 |
| YZH_070 | h- ade6-M216 ura4-D18 leu1-32 hrp3-13xmyc-kanMX6 |
| YZH_276 | h- ade6-M216 ura4-D18 leu1-32 ura4-D18Δ::kanMX6-Pr_Spombe_Adh1-Ura4 |
| YZH_278 | h- ade6-M216 ura4-D18 leu1-32 ura4-D18Δ::kanMX6-Pr_AT1TE70815-Sp_Ura4 |
| YJY_023 | h+ ade6-M216 ura4-D18 leu1-32 fft1Δ::hygMX |
| YJY_024 | h+ ade6-M216 ura4-D18 leu1-32 fft2Δ::kanMX |
| YJY_025 | h+ ade6-M216 ura4-D18 leu1-32 fft3Δ::kanMX |
| YJY_026 | h+ ade6-M216 ura4-D18 leu1-32 hrp1Δ::kanMX |
| YJY_027 | h+ ade6-M216 ura4-D18 leu1-32 hrp3Δ::kanMX |
| YJY_028 | h+ ade6-M216 ura4-D18 leu1-32 mit1Δ::kanMX |
| YJY_035 | h+ ade6-M216 ura4-D18 leu1-32 rrp1Δ::kanMX |
| YJY_036 | h+ ade6-M216 ura4-D18 leu1-32 rrp2Δ::kanMX |
| YJY_037 | h+ ade6-M216 ura4-D18 leu1-32 snf22Δ::hygMX |
| YJY_038 | h+ ade6-M216 ura4-D18 leu1-32 irc20Δ::kanMX |
| YJY_039 | h+ ade6-M216 ura4-D18 leu1-32 swr1Δ::kanMX |
| YJY_061 | h+ ade6-M216 ura4-D18 leu1-32 hrp1Δ::kanMX hrp3Δ |
| YJY_064 | h+ ade6-M216 ura4-D18 leu1-32 hrp1Δ::hrp1-CD(hrp3) hrp3Δ |
| YJY_065 | h+ ade6-M216 ura4-D18 leu1-32 hrp1Δ::kanMX hrp3Δ::hrp3-CD(hrp1) |
| YJY_070 | h+ ade6-M216 ura4-D18 leu1-32 hrp1Δ::hrp1-CT(hrp3) hrp3Δ |
| YJY_071 | h+ ade6-M216 ura4-D18 leu1-32 hrp1Δ::kanMX hrp3Δ::hrp3-CT(hrp1) |
| YJY_095 | h- ade6-M216 ura4-D18 leu1-32 ura4-D18Δ::kanMX6-Pr_rap1(As)-Sp_Ura4 |
| YJY_096 | h- ade6-M216 ura4-D18 leu1-32 ura4-D18Δ::kanMX6-Pr_atg9(As)-Sp_Ura4 |
| YJY_097 | h- ade6-M216 ura4-D18 leu1-32 ura4-D18Δ::kanMX6-Pr_crt10(As)-Sp_Ura4 |
| YJY_098 | h- ade6-M216 ura4-D18 leu1-32 ura4-D18Δ::kanMX6-Pr_spb70(As)-Sp_Ura4 |
| YJY_099 | h- ade6-M216 ura4-D18 leu1-32 ura4-D18Δ::kanMX6-Pr_orc4(As)-Sp_Ura4 |
| YZH_276 | h- ade6-M216 ura4-D18 leu1-32 ura4-D18Δ::kanMX6-Pr_Spombe_Adh1-Ura4 |
| YZH_278 | h- ade6-M216 ura4-D18 leu1-32 ura4-D18Δ::kanMX6-Pr_AT1TE70815-Sp_Ura4 |
| YJY_122 | h+ ade6-M216 ura4-D18 leu1-32 prf1Δ |
| YJY_154 | h+ ade6-M216 ura4-D18 leu1-32 hrp1Δ::kanMX hrp3Δ prf1Δ |
| YJY_177 | h- ade6-M216 ura4-D18 leu1-32 hrp3-CHCT_R1(AlaScan)-13xmyc-kanMX |
| YJY_178 | h- ade6-M216 ura4-D18 leu1-32 hrp3-CHCT_R2(AlaScan)-13xmyc-kanMX |
| YJY_179 | h- ade6-M216 ura4-D18 leu1-32 hrp3-CHCT_R3(AlaScan)-13xmyc-kanMX |
| YJY_180 | h- ade6-M216 ura4-D18 leu1-32 hrp3-CHCT_R4(AlaScan)-13xmyc-kanMX |
| YJY_181 | h- ade6-M216 ura4-D18 leu1-32 hrp3-CHCTΔ-13xmyc-kanMX |
